# Transcriptional regulation of the TASK-1 potassium channel by ETV1 – Implications for atrial excitability

**DOI:** 10.64898/2026.03.12.711402

**Authors:** Moritz Beck, Felix Wiedmann, Manuel Kraft, Patrick Laurette, Amelie Paasche, Lanzer Jan, Max Jamros, Ciara Malchin, Paul Henri Ziehmer, Christian Goetz, Marcin Zaradzki, Rawa Arif, Matthias Karck, Norbert Frey, Ralf Gilsbach, Constanze Schmidt

**Author notes:** Correspondence: Prof. Constanze Schmidt, MD, FESC, FEHRA University Medical Center Goettingen Department of Cardiology and Pneumology Robert-Koch-Str. 40, D-37075 Goettingen, Germany. Authors contributed equally (Moritz Beck, Felix Wiedmann, Manuel Kraft).

## Abstract

**Background:** Atrial fibrillation (AF), the most common sustained arrhythmia, is driven by electrical and structural remodelling, including altered ion channel expression. The atrial-specific potassium channel TASK-1 regulates action potential duration (APD) and is differentially expressed in AF and left ventricular dysfunction, but the mechanisms controlling its expression are not well understood.

**Objective:** This study examines whether the transcription factor ETV1 regulates TASK-1 and contributes to atrial electrical remodelling.

**Methods:** Atrial tissue from patients with and without AF was analysed to assess the relationship between ETV1 and TASK-1 (*KCNK3*) expression. In HL-1 cardiomyocyte-like cells and native fibroblasts, ETV1 activity was reduced using pharmacological inhibition or siRNA-mediated knockdown. TASK-1 expression, TASK-1 current, and APD at 90% repolarization were measured. Pacing experiments tested activity-dependent TASK-1 regulation. Direct transcriptional regulation was evaluated using ChIP-qPCR and ChIP-seq to detect ETV1 binding at the *KCNK3* promoter.

**Results:** ETV1 and TASK-1 levels were positively correlated in human atrial tissue. In HL-1 cells and fibroblasts, ETV1 inhibition or knockdown decreased TASK-1 expression and current and selectively prolonged APD90. Pacing-induced upregulation of TASK-1 was prevented by ETV1 inhibition, indicating a protective effect against pro-arrhythmic remodelling. ChIP-qPCR and ChIP-seq confirmed direct ETV1 binding to the *KCNK3* promoter.

**Conclusion:** ETV1 directly regulates TASK-1 expression and contributes to atrial electrical remodelling, identifying ETV1 as a potential upstream therapeutic target in AF.

**Graphical abstract:** 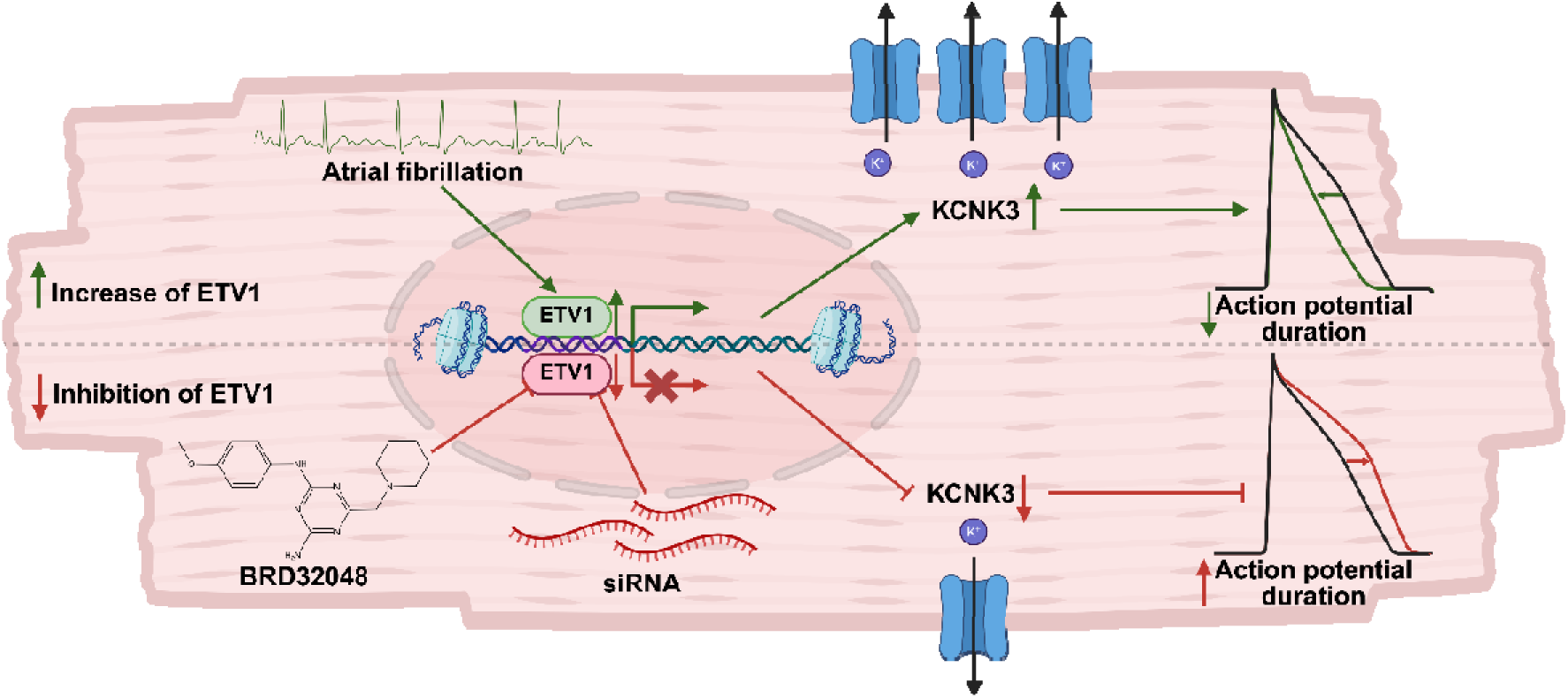

**Translational perspective:** Atrial fibrillation is sustained by maladaptive electrical remodelling that remains insufficiently addressed by current rhythm-control therapies. Direct inhibition of individual ion channels has shown efficacy but is limited by phenotype dependence and proarrhythmic risk.

The present data identify ETV1 as an upstream transcriptional regulator of the atrial-specific potassium channel TASK-1. Modulation of ETV1 reduced TASK-1 expression, prolonged atrial repolarisation, and prevented tachycardia-induced electrical remodelling in vitro.

Targeting ETV1 may therefore represent a disease-modifying strategy that intervenes earlier in the remodelling cascade than conventional antiarrhythmic drugs. This approach could enable phenotype-guided therapy in atrial cardiomyopathy, particularly in patients with preserved ventricular function, and warrants validation in translational large-animal and clinical studies.

## Introduction

Atrial fibrillation (AF) is the most common sustained cardiac arrhythmia, characterized by disorganized atrial electrical activity and loss of coordinated atrial contraction. It is associated with an increased risk of stroke, acute heart failure and cardiovascular morbidity ^1^. The pathophysiology of AF involves progressive structural and electrical remodelling processes that increase the susceptibility to arrhythmogenesis and promote arrhythmia maintenance ^2^. While structural remodelling implies fibrosis and architectural changes, electrical remodelling encompasses abnormal regulation of ion channels and conduction-related proteins ^3–5^.

Among the ion channels involved in AF-associated electrical remodelling, the two-pore domain potassium channel (K_2P_) TASK-1 has emerged as one of the key players ^6^. In the human heart, TASK-1 is predominantly expressed in the atria and significantly upregulated in patients with AF, thereby contributing to the characteristic shortening of the atrial action potential duration (APD) ^6^. This APD shortening promotes re-entrant circuits, which help sustain AF ^3^. Inhibition of TASK-1 has been shown to prolong the APD to levels observed in sinus rhythm (SR), thereby leading to the termination of AF ^6,7^. In contrast, this effect is attenuated in patients with reduced left ventricular function (LVF), as TASK-1 expression is downregulated in these individuals ^8^. The antiarrhythmic effect of TASK-1 inhibition was demonstrated in a preclinical large animal model of AF using both pharmacological modulation and AAV9-based gene therapy ^7,9,10^. A clinical first-in-man study on the use of a TASK-1 inhibitor, the DOCTOS trial (EudraCT No. 2018-002979-17), DOxapram Conversion TO Sinus rhythm study, is near publication. Despite these insights, the upstream regulatory mechanisms controlling atrial TASK-1 expression remain poorly understood ^4,7^.

Transcription factors are central in orchestrating cardiac gene expression during development and in disease states, including hypertrophy and arrhythmias ^11^. Among these, E twenty-six (ETS) translocation variant 1 (ETV1) is abundantly expressed in the heart, particularly within the conduction system ^12,13^. ETV1 has further been implicated in atrial remodelling processes in murine disease models, and preliminary evidence suggests that it may regulate TASK-1 expression ^14^.

Therefore, the aim of this study was to investigate the role of ETV1 in regulating atrial TASK-1 expression.

## Methods

### Human Tissue Samples

Human atrial tissue samples were obtained from patients undergoing open-heart surgery after written informed consent and approval by the local ethics committee (Medical Faculty Heidelberg, Germany; approval S-017/2013). Samples from patients in sinus rhythm and atrial fibrillation were used for electrophysiological, molecular, and transcriptomic analyses. Tissue handling and storage were standardized. Detailed patient characteristics, sample numbers per experiment, and processing protocols are provided in the Supplementary Methods and Tables S1–S3.

### Animal Experiments

All animal experiments were approved by the local authorities and conducted in accordance with NIH guidelines, EU Directive 2010/63/EU, and German animal protection law. Experiments were performed in pigs (*Sus scrofa domesticus*) and *Xenopus laevis*. A chronic porcine AF model was established by atrioventricular nodal ablation, pacemaker implantation, and repeated atrial burst pacing. Tissue harvesting was performed under deep anaesthesia. Full anaesthesia protocols, pacing parameters, and animal numbers are described in the Supplementary Methods.

### Cell Isolation and Cell Culture

Atrial cardiomyocytes and fibroblasts were isolated from human and porcine atrial tissue using established enzymatic protocols. HL-1 cardiomyocytes and HEK293T cells were cultured under standard conditions. Details on isolation procedures, culture media, passage numbers, and experimental treatments are provided in the Supplementary Methods.

### Electrophysiology

Whole-cell patch-clamp recordings were performed on freshly isolated atrial cardiomyocytes and HL-1 cells to assess potassium currents and action potentials. Two-electrode voltage-clamp recordings were performed in *Xenopus laevis* oocytes expressing human TASK-1 channels. Recording solutions, stimulation protocols, and analysis methods are described in detail in the Supplementary Methods.

### Statistical Analysis

Data are presented as mean ± SEM unless stated otherwise. Appropriate parametric statistical tests were applied as indicated. A two-sided p-value < 0.05 was considered statistically significant. Full statistical methods are provided in the Supplementary Methods.

## Results

### TASK-1 and ETV1 are Co-regulated in Response to Rhythm Status and Left Ventricular Function

AF is a heterogeneous clinical condition characterized by different underlying subtypes of atrial cardiomyopathy, driven by mechanisms such as fibrosis, extracellular matrix remodelling and dysregulated ion channel expression (Figure 1A). Among the ion channels implicated in electrical remodelling, the atrial-specific potassium channel TASK-1 has emerged as a key contributor. However, the molecular mechanisms controlling its expression remain largely elusive.

**Figure 1:**
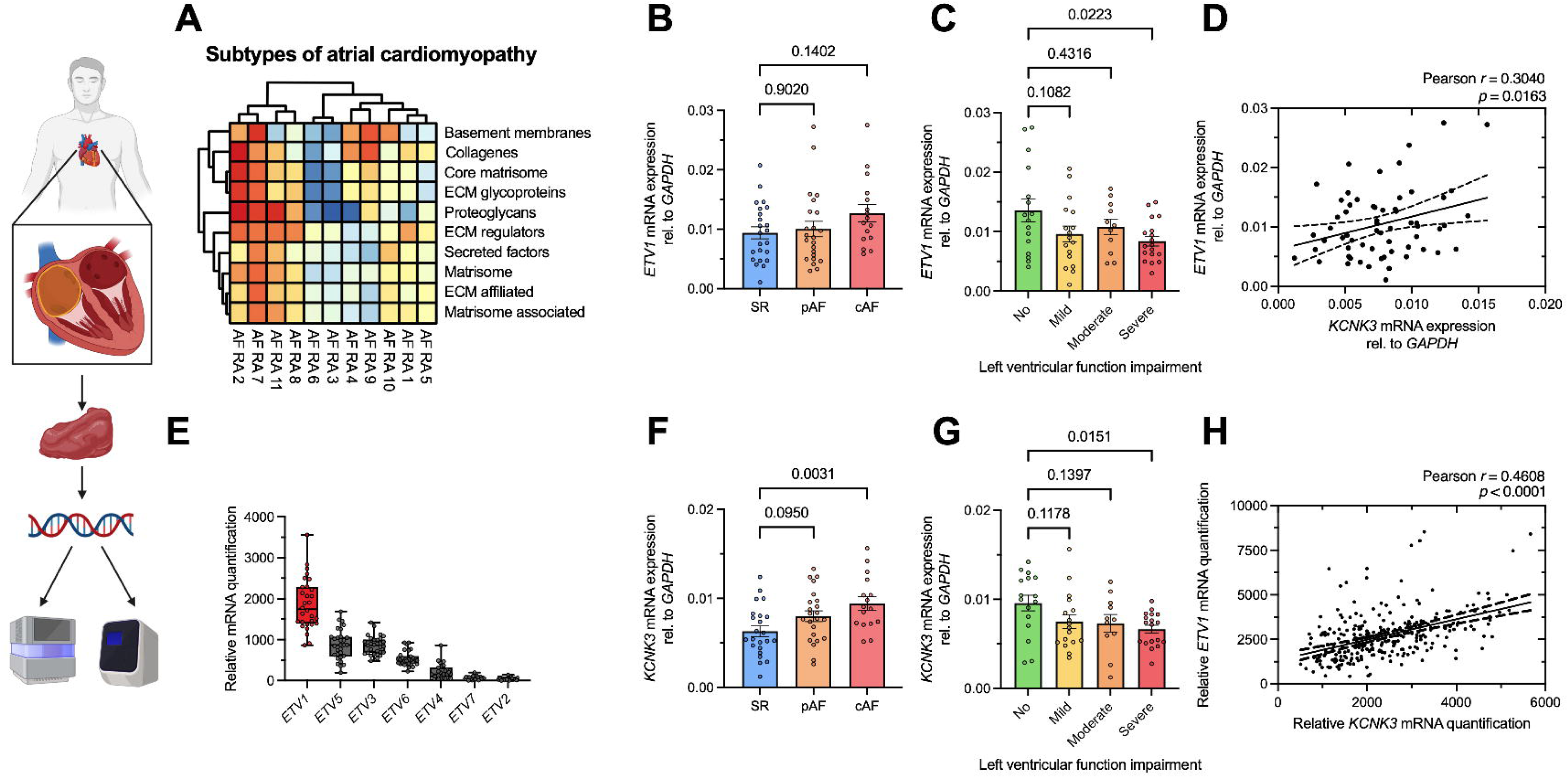
Correlation of TASK-1 and ETV1 expression. *Left*: Schematic overview of the workflow from sample collection to RNA isolation, DNA preparation and quantification by real-time quantitative polymerase chain reaction (RT-qPCR) or RNA sequencing (RNAseq). **A**: Cluster analysis of right atrial (RA) appendage tissue samples from *n* = 11 atrial fibrillation (AF) patients revealing distinct molecular subtypes of atrial cardiomyopathy. Samples were clustered based on RNAseq results of fibrosis- and extracellular matrix (ECM)-related markers. **B–H**: Expression levels of *KCNK3* (encoding for TASK-1) and *ETV1* are associated with atrial rhythm (B, F) and left ventricular function (LVF, C, G). **B–C**: qPCR analysis of *ETV1* mRNA levels, stratified by atrial rhythm (B): sinus rhythm (SR, *n* = 23), paroxysmal AF (pAF, *n* = 23), and chronic AF (cAF, *n* = 16) or by LVF (C): normal (no, *n* = 16), mildly (mild, *n* = 17), moderately (moderate, *n* = 11), and severely (severe, *n* = 18) reduced LVF. **D**: Correlation analysis of *KCNK3* and *ETV1* mRNA levels quantified using real time quantitative polymerase chain reaction in the cohort shown in panel B–C. **E**: Relative quantification of *ETV1*–*7* mRNA levels from RNAseq of RA appendage samples (*n* = 30; SR: *n* = 15, AF: *n* = 15), analysed using the DEseq2 algorithm. **F–G**: Quantification of *KCNK3* mRNA levels by qPCR in right atrial appendage samples stratified by atrial rhythm (F) and degree of left ventricular dysfunction (G), corresponding to the cohorts shown in panels B–D. **H**: Correlation of *KCNK3* and *ETV1* mRNA expression levels based on previously published RNA sequencing data from 276 human left atrial appendage samples, as reported by Tchou *et al.*, Gene Expression Omnibus (GEO) accession number GSE69890 ^15^. Data are presented as mean ± standard error of the mean (SEM). Group comparisons were performed using ordinary one-way ANOVA with Dunnett’s correction for multiple comparisons and *p*-values < 0.05 were considered statistically significant. For correlation analysis, Pearson correlation coefficients and respective *p*-values are provided as insets.

To explore potential shared regulatory patterns, *ETV1* mRNA expression was quantified by real-time quantitative polymerase chain reaction (RT-qPCR) in human atrial appendage samples from patients in SR, paroxysmal AF (pAF) and chronic AF (cAF; Figure 1B–H). A slight, non-significant increase in *ETV1* expression was observed in pAF compared to SR (*p* = 0.90; *n* = 23; Figure 1B). This trend was even more pronounced in cAF samples, though statistical significance was not reached (*p* = 0.14; *n* = 16–23; Figure 1B). Stratification by LVF revealed a trend toward decreased *ETV1* expression with impairment of left ventricular dysfunction. Patients with preserved LVF exhibited highest *ETV1* mRNA levels, while those with severely reduced LVF showed the lowest expression (no impairment vs. mild: *p* = 0.11, *n* = 16–17; no impairment vs. moderate: *p* = 0.43, *n* = 11–16; no impairment vs. severe: *p* = 0.022, *n* = 16–18; Figure 1C).

A similar expression pattern for *KCNK3* mRNA, encoding for TASK-1 channel subunits, was observed in this patient cohort: *KCNK3* mRNA levels were elevated in the pAF group compared to SR, but this difference did not reach statistical significance (*p* = 0.095; *n* = 23; Figure 1F). In contrast, a significant increase in *KCNK3* mRNA levels was observed in cAF samples relative to SR (*p* = 0.003; *n* = 16–23; Figure 1F). Stratification by LVF revealed a pattern similar to that of *ETV1*, with the highest *KCNK3* mRNA expression levels in patients with preserved LVF and a stepwise decline across mild, moderate and severe impairment (no impairment vs. mild: *p* = 0.12, *n* = 16–17; no impairment vs. moderate: *p* = 0.14, *n* = 11–16; no impairment vs. severe: *p* = 0.015, *n* = 16–18; Figure 1G). Correlation analysis of *ETV1* and *KCNK3* mRNA expression in this cohort of human right atrial appendage samples revealed a moderate positive correlation (Pearson *r* = 0.3040, p = 0.016; Figure 1D). This association was further supported by RNA sequencing data from 276 human left atrial appendage samples, previously published by Tchou et al. (GEO accession GSE69890) ^15^, which showed a stronger positive correlation between relative *ETV1* and *KCNK3* mRNA counts (Pearson *r* = 0.4608, *p* < 0.0001; Figure 1H). These parallel expression patterns of *ETV1* and *KCNK3* suggest a potential transcriptional association between the two.

Furthermore, mRNA expression levels of the entire ETV transcription factor family were compared by RNA sequencing analysis (DEseq2) in a cohort of 30 patient samples equally divided between SR and AF. *ETV1* emerged as the most abundantly expressed member in atrial tissue, highlighting its potential relevance and supporting the focus on this transcription factor for further analysis (Figure 1E).

To extend the analysis to the entire ETV transcription factor family, RT-qPCR was performed on the same cohort of patients analysed for *ETV1* and *KCNK3*. AF-dependent regulation patterns were observed for *ETV2*, *ETV5*, and *ETV6* (Figure S1A). These findings were further supported by RNA sequencing data from Figure 1E when stratified by rhythm status (Figure S2). Stratification by LV dysfunction revealed a significant downregulation of *ETV4* and upregulation of *ETV6* in patients with severe LVF impairment (Figure S1B). Finally, correlation analysis showed a positive association of *KCNK3* with *ETV5* and *ETV6*, and a negative association with *ETV2* (Figure S1C).

Taken together, these results suggest a complex and differential regulation of ETV family members in the atrium. Among them, *ETV1* emerged as the most consistently expressed and strongly correlated with *KCNK3*, supporting its role as a key candidate for further investigation in the context of AF pathophysiology.

### Comparable Regulation in Fibroblasts Across Species

Human atrial fibroblasts isolated from AF patient tissue samples exhibited significantly higher *ETV1* mRNA expression compared to those from SR patients (*p* = 0.0003; *n* = 3–6; Figure S3A). Treatment with the ETV1 inhibitor BRD32048 led to a more pronounced reduction of *ETV1* expression in AF-derived fibroblasts (−70.98%; *p* < 0.0001; *n* = 3–6) than in those from SR patients (−57.76%; *p* < 0.0001; *n* = 3–6; Figure S3A). A similar trend was observed for *KCNK3* mRNA (Figure S3B), which showed elevated expression levels in AF-derived fibroblasts relative to SR (*p* = 0.057; *n* = 3–6), and a greater reduction upon BRD32048 treatment in the AF group (−53.72% vs. −41.66%; AF: *p* = 0.032; *n* = 3–6). Notably, *ETV1* mRNA levels were substantially lower in fibroblasts than in cardiomyocytes (*p* < 0.001; *n* = 7–8; Figure S3C), underscoring a cell-type-specific role of ETV1.

Finally, these observations were recapitulated in porcine atrial fibroblasts (Figure S3D–E). While *ETV1* mRNA expression did not differ significantly between fibroblasts from pigs in SR and those with artificially induced AF via right atrial burst-stimulation (*p* = 0.38; *n* = 3–6; Figure S3D), *KCNK3* mRNA levels were markedly increased in AF fibroblasts (*p* < 0.0001; *n* = 3–6; Figure S3E). Treatment with BRD32048 resulted in a substantial reduction of both *ETV1* (*p* < 0.0001; *n* = 6; Figure S3D) and *KCNK3* mRNA (*p* < 0.0001; *n* = 6; Figure S3E) expression in AF fibroblasts. In contrast, SR fibroblasts showed a pronounced decrease in *ETV1* mRNA levels (*p* = 0.0001; *n* = 3–6), whereas *KCNK3* mRNA levels expression remained largely unchanged (*p* = 0.55; *n* = 3–6). Consistent with the human data, *KCNK3* mRNA levels were significantly higher in cardiomyocytes compared to fibroblasts (*p* = 0.003; *n* = 10–11; Figure S3F), further supporting a distinct functional role of TASK-1 in cardiomyocytes.

### Functional and Posttranscriptional ETV1 Inhibition Leads to Reduced TASK-1 Expression

To investigate the functional relationship between ETV1 and TASK-1, pharmacological inhibition and siRNA-mediated knockdown of ETV1 were performed in HL-1 murine atrial cardiomyocyte-like cells. Treatment of HL-1 cells with the pharmacological ETV1 inhibitor BRD32048 a small molecule disrupting ETV1-p300 interaction and thereby promoting its degradation ^16^, did not significantly alter *Etv1* mRNA expression over a 72-hour period (*p* = 0.49; *n* = 6; Figure 2A), indicating that the compound neither promotes *Etv1* mRNA degradation nor induces compensatory transcriptional upregulation. However, *Kcnk3* mRNA expression was markedly reduced over the 72-h treatment period (control vs. 48 h: *p* = 0.009; *n* = 6; control vs. 72 h: *p* = 0.009; *n* = 6; Figure 2B). This effect was time-dependent, reaching statistical significance at 48 h and 72 h, with an overall reduction of nearly 50% by 72 h. A corresponding decrease was observed at the protein level, with a modest trend at 24 h that became significant after 48 h (control vs. 48 h: *p* = 0.009; *n* = 3; control vs. 72 h: *p* = 0.002; *n* = 3; Figure 2C).

**Figure 2:**
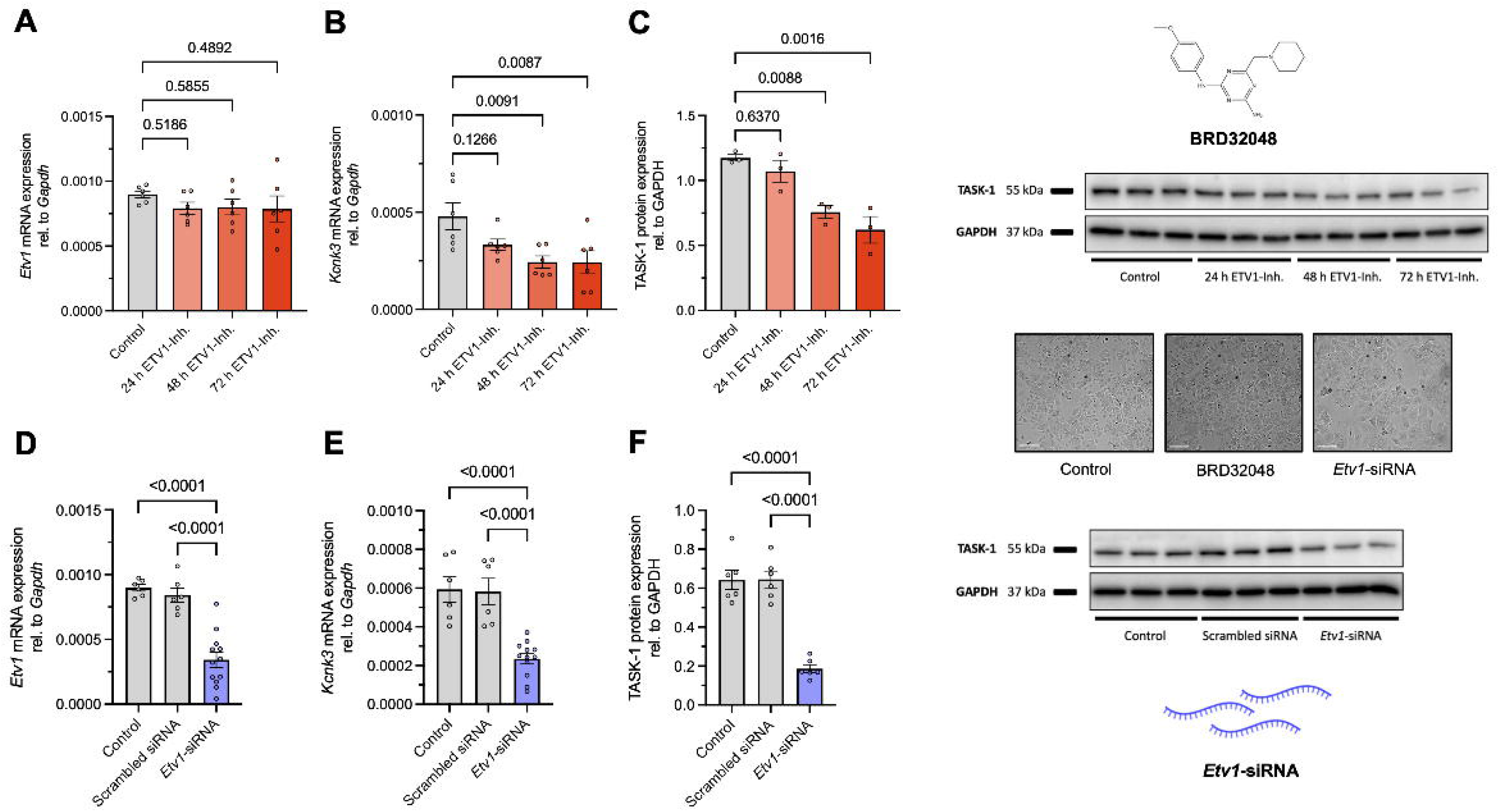
ETV1 inhibition leads to reduced TASK-1 expression. A–B: *Etv1* mRNA (A, *n* = 6) and *Kcnk3* mRNA (B, *n* = 6; encoding for TASK-1) expression relative to *Gapdh* following treatment of HL-1 cells with 10 µM of the specific ETV1 inhibitor BRD32048 for 24 h, 48 h, and 72 h. **C**: *Left*: TASK-1 protein expression (*n* = 3) relative to GAPD under the same treatment conditions. *Right*: Corresponding immunoblot. **D–E**: *Etv1* (D, *n* = 6–12) and *Kcnk3* mRNA (E, *n* = 6–12) expression relative to *Gapdh* in HLL1 cells 48 h after transfection with *Etv1*-targeting siRNA or scrambled control. **F**: *Left*: TASK-1 protein expression (*n* = 6) relative to GAPDH in HL-1 cells 48 h after transfection with *Etv1*-targeting siRNA or scrambled control. *Right*: Representative immunoblot. Data are presented as mean ± standard error of the mean (SEM). Scalebar depicts 50 µm. Ordinary one-way ANOVA was used for group comparisons with Dunnett’s correction (A–C) and Šídák’s correction (D–F) for multiple comparisons. Statistical significance was defined as *p*-values < 0.05.

Transfection with siRNA targeting *Etv1* resulted in a robust and statistically significant reduction in *Etv1* mRNA levels (*p* < 0.0001; *n* = 6–12; Figure 2D) corresponding to an approximate 60% decrease and indicating a direct transcript degradation by the siRNA. Scrambled siRNA had no effect on *Etv1* expression. Further, a significant downregulation of *Kcnk3* mRNA could be detected at 48 h post-transfection (p < 0.0001; *n* = 6–12; Figure 2E). This reduction was also mirrored at the protein level, with TASK-1 reduced by 70% upon *Etv1* knockdown compared to control and scrambled groups (*p* < 0.0001; Figure 2F).

### ETV1 Inhibition Prolongs APD**□□** and Suppresses TASK-1 Currents in HL-1 cells

Incubation of HL-1 cells with the ETV inhibitor BRD32048 for up to 72 h did not result in significant changes in APD at 50% repolarisation (APD_50_), although a non-significant trend towards prolongation was observed (Control vs. 72 h: *p* = 0.54; *n* = 15–20; Figure 3A–B). In contrast, a significant prolongation was observed in APD at 90% repolarisation (APD_90_), consistent with the previously described reduced TASK-1 expression (Control vs. 48 h: *p* = 0.046, *n* = 15–16; Control vs. 72 h: *p* = 0.022, *n* = 15–20; Figure 3B). This finding was further supported by a time-dependent reduction in TASK-1 current, isolated via application of the high-affinity TASK-1 inhibitor A293 (Control vs. 48 h: *p* = 0.009, *n* = 7–12; Control vs. 72 h: *p* = 0.003, *n* = 8–12; Figure 3C–D). No significant changes were observed in action potential amplitude (APA) or resting membrane potential (RMP, Figure 3B).

**Figure 3:**
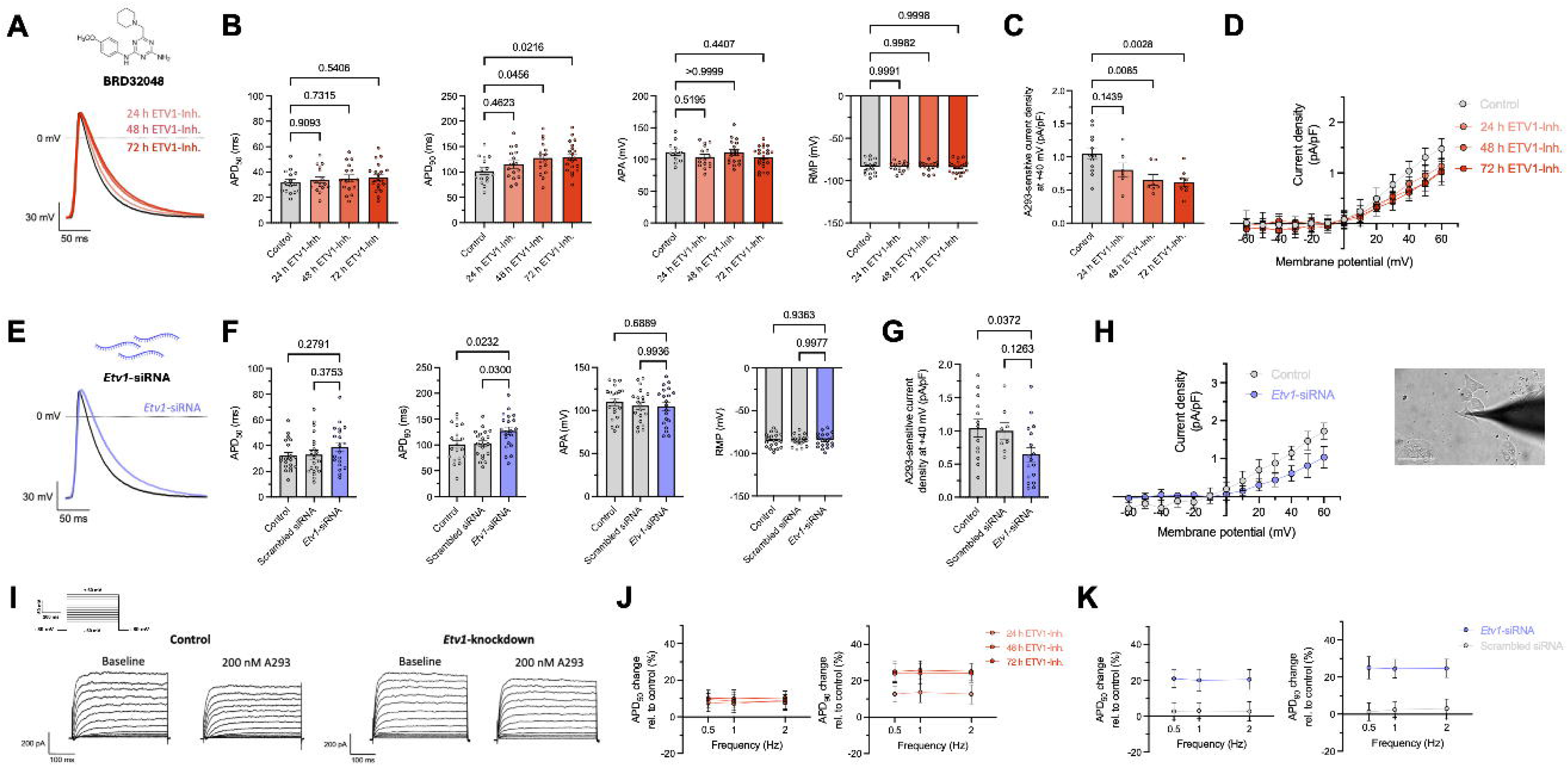
Effects of ETV1 inhibition on action potential (AP) parameters and TASK-1 currents. **A**: Representative APs, recorded from HL-1 cells by patch-clamp measurements after 24 h, 48 h and 72 h treatment with 10 µM of the specific ETV1 inhibitor BRD32048 (ETV1-Inh.) compared to untreated (Control) cells. **B**: Action potential duration (APD) at 50% and 90% of repolarization (APD_50_, APD_90_), action potential amplitude (APA), and resting membrane potential (RMP) in HL-1 cells treated with BRD32048 (10 µM; ETV1-Inh.) or untreated (Control; *n* = 15–20). **C**: Functional TASK-1 current densities (isolated using the high-affinity TASK-1 inhibitor A293, 200 nM) as A293-sensitive current density in HL-1 cells after 24 h, 48 h, and 72 h of BRD32048 treatment (10 µM; ETV1-Inh.) compared to untreated cells (Control; *n* = 7–12). **D**: Current-voltage relationship of A293-sensitive current densities of HL-1 cells under the same conditions as in (C) **E**: Representative action potential recordings from HL-1 cells, obtained by patch-clamp measurements 48 h after transfection with *Etv1*-siRNA, scrambled siRNA, or in untreated controls (*n* = 8–17). **F**: APD_50_, APD_90_, APA, and RMP in HL-1 cells 48 h after transfection with *Etv1*-siRNA, scrambled siRNA (Scrambled siRNA) or no transfection (Control; *n* = 20). **G**: A293-sensitive current density of HL-1 cells 48 h post-transfection with *Etv1*-siRNA, scrambled siRNA, or in untreated controls (Control; *n* = 8–17). **H**: Current-voltage-relationship of A293-sensitive current density of HL-1 cells under the same conditions as in (G). **I**: Representative current traces recorded from untreated (Control) HL-1 cells and cells with ETV1 knockdown by application of the depicted pulse protocol, both under baseline conditions and after exposure to 200 nM A293. **J–K**: Frequency dependence of change in APD_50_ and APD_90_ of HL-1 cells treated with BRD32048 for 24 h, 48 h and 72 h (*n* = 4–6; J) and in cells transfected with *Etv1-*or scrambled siRNA (*n* = 5–6; K). Data are presented as mean ± standard error of the mean (SEM). Ordinary one-way ANOVA was used for group comparison with Dunnett’s correction (B–C) and Šídák’s correction (F–G) for multiple testing. Statistical significance was defined as *p*-values < 0.05. Zero current and potential levels are indicated by dashed lines. Pulse protocols and scale bars are shown as insets. Scalebar depicts 20 µm.

Transfection of HL-1 cells with siRNA targeting *Etv1* recapitulated the effects observed with pharmacological treatment. While APD_50_ showed a non-significant trend towards prolongation (*p* = 0.28; *n* =20), APD_90_ was significantly increased (*p* = 0.023; *n* = 20; Figure 3E–F). A293-sensitive current was also significantly reduced following siRNA treatment (*p* = 0.037; *n* = 13–17; Figure 3G–I), confirming decreased TASK-1 function. As with pharmacological ETV1 inhibition, neither APA nor RMP were affected (Figure 3F). Finally, the prolongation of APD_90_ following ETV-1 inhibition – whether pharmacological or siRNA-mediated – was consistent across varying stimulation frequencies (Figure 3 J–K).

Taken together, these results demonstrate that ETV1 inhibition reduces TASK-1 current via downregulation of TASK-1 expression, leading to a selective prolongation of APD_90_ with minimal impact on other AP parameters.

To evaluate whether BRD32048 directly blocks the TASK-1 channel pore, *Xenopus laevis* oocytes injected with cRNA encoding human TASK-1 were acutely superfused with 10 µM BRD32048 for 30 minutes, while TASK-1 currents were recorded every 2 minutes. No changes in current amplitude were observed during the treatment period, indicating that BRD32048 does not exert acute, pore-blocking effects on the channel. These findings support the conclusion that the observed reductions in TASK-1 currents are mediated indirectly through ETV1 inhibition rather than by direct interaction with the channel pore (Figure S4).

### Modest Reduction of TASK-1 Currents After Short-Term ETV1 Inhibition

Freshly isolated human atrial cardiomyocytes were incubated with 1 µM BRD32048 for 6 h, followed by patch-clamp measurements to assess A293-sensitive current as surrogate for TASK-1.

In untreated control cells, TASK-1 current was highest in cardiomyocytes from patients with cAF (1.84 pA/pF; Figure 4C), followed by those from patients with pAF (0.99 pA/pF; Figure 4B) and SR (0.41 pA/pF; Figure 4A). Following BRD32048 treatment, a modest trend towards a reduced TASK-1 current was observed across all groups (SR: −18.7 %, pAF: −14.5 %, cAF: −11.2 %; Figure 4A-C), although these changes did not reach statistical significance. This limited effect is likely attributable to the short incubation period and the expected delay of transcriptional responses.

**Figure 4:**
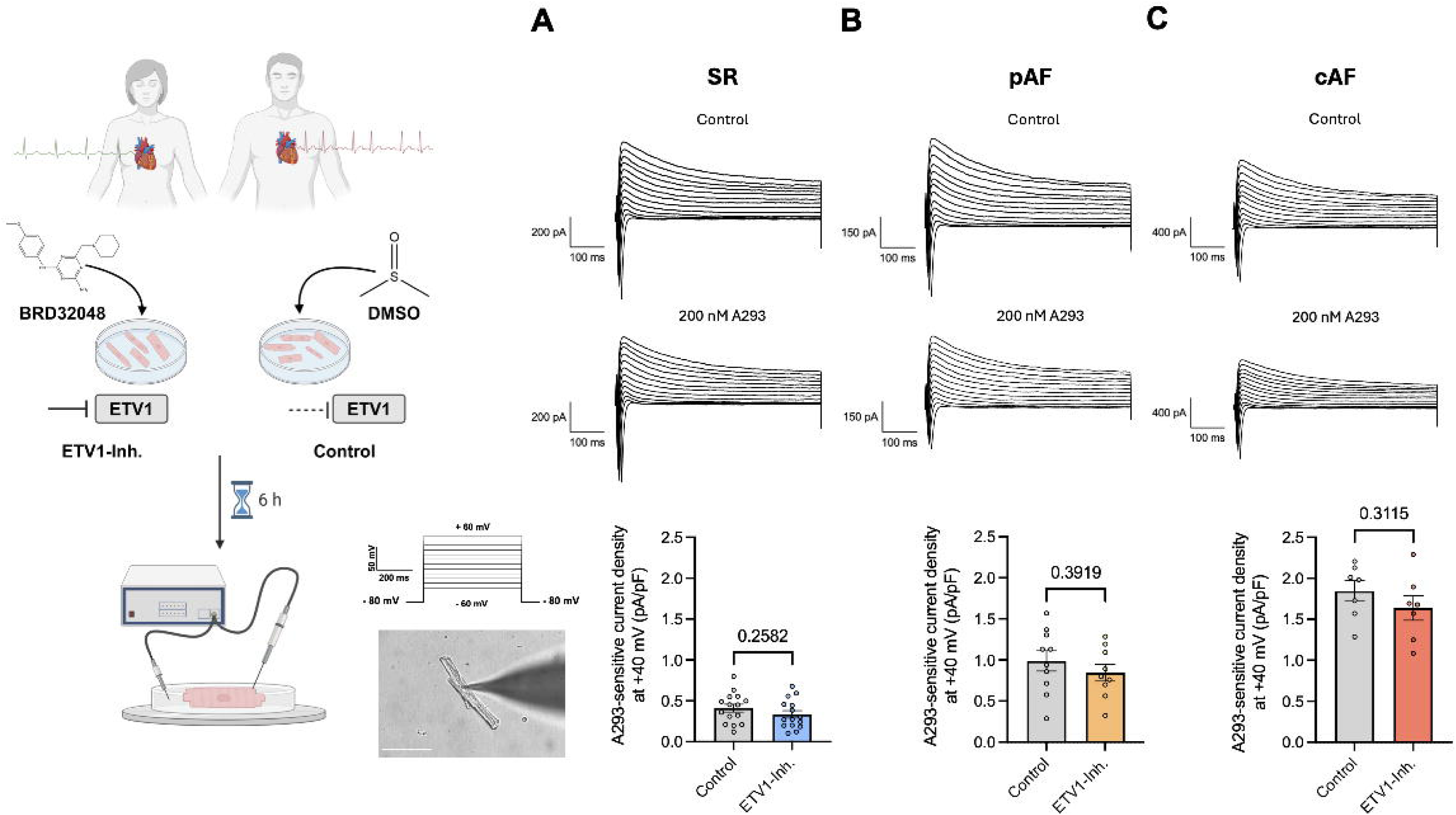
Effects of ETV1 inhibition on the TASK-1 current in human atrial cardiomyocytes. **A–C:** Freshly isolated cardiomyocytes from patient-derived tissue samples were incubated for 4–6 h with the specific ETV1 inhibitor BRD32048 (10 µM). The A293-sensitive current was measured in cardiomyocytes from patients in sinus rhythm (SR, A, *n* = 15 cardiomyocytes, isolated from N = 5 individual patients), paroxysmal atrial fibrillation (pAF, B, *n/N* = 9–10/4), and chronic atrial fibrillation (cAF, C, *n/N* = 7/3). Outward potassium currents were quantified at the end of the +40 mV pulse and are presented as mean ± standard error of the mean (SEM). Representative current traces are shown at the top, traces after incubation with 200 nM A293 in the middle and corresponding quantification at the bottom. A schematic overview of cardiomyocyte isolation and patch-clamp recording is shown on the left. The pulse protocol and scalebars are provided as insets. Zero current levels are indicated by dotted lines. Unpaired Student’s *t*-tests were used to compare treatment versus control groups, where *p*-values < 0.05 were considered statistically significant. Scalebar depicts 75 µm.

### ETV1 Inhibition Protects Against Tachypacing Induced Electrical Remodelling *in vitro*

Atrial cardiomyopathy encompasses distinct disease subtypes that appear to arise in response to specific clinical stressors. Translational evidence suggests that these stressors critically influence the pattern of atrial remodelling. To explore this experimentally, various stress conditions including angiotensin II, isoprenaline, hypoxia, hyperglycaemia and tachypacing were modelled *in vitro* using cultured HL-1 cells, and their impact on ETV1 and TASK-1 expression was examined (Figure 5 and Figure S5). Among all tested models, tachypacing via field stimulation proved to be the modality that most effectively reproduced the molecular and electrical phenotype of TASK-1–based electrical remodelling (Figure 5 and Figure S5), prompting us to focus for subsequent experiments on this *in vitro* model.

**Figure 5:**
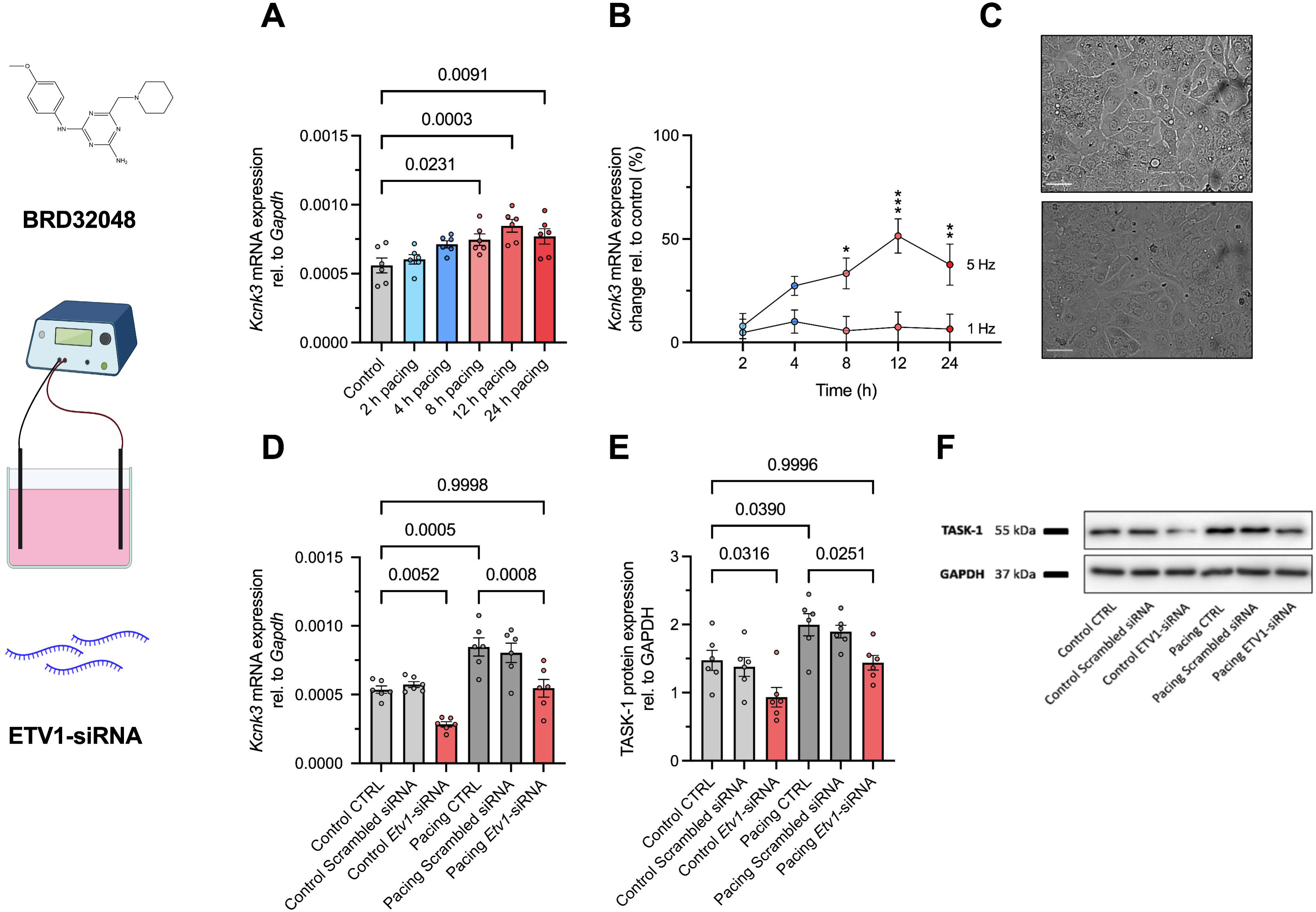
ETV1 inhibition prevents tachypacing-induced TASK-1 dysregulation in HL-1 cells. A–B: HL-1 cells were subjected to field stimulation with 1 Hz (Control) or 5 Hz for 2 h, 4 h, 8 h, 12 h, and 24 h. *Kcnk3* mRNA was quantified relative to *Gapdh* and shown as absolute values (A) and values normalized to the 1 Hz control (B; *n* = 6). **C**: Representative images of HL-1 cells before (top) and after 24 h of stimulation (bottom). Scalebar depicts 50 µm. **D–F**: HL-1 cells transfected with *Etv1* targeting siRNA, scrambled siRNA, or untransfected controls (CTRL) were paced using field stimulation at 5 Hz (Pacing) and 1 Hz (Control) for 12 h. **D–E**: *Kcnk3* mRNA (D) and TASK-1 protein expression (E) relative to GAPDH (*n* = 6). **F**: Representative immunoblot showing *TASK-1* protein expression. Data are shown as mean ± standard error of the mean (SEM). Group comparisons were performed using ordinary one-way ANOVA with Šídák’s correction for multiple comparisons. Statistical significance was defined as *p*-values < 0.05.

Field stimulation of HL-1 cells led to a significant upregulation of *Kcnk3* mRNA levels, evident after 8 h (*p* = 0.023; *n* = 6) and peaking at 12 h (*p* = 0.0003; *n* = 6; Figure 5A). Comparing stimulation at 1 Hz and 5 Hz revealed that the increase in Kcnk3 mRNA levels occurred only at the high frequency (Figure 5B), indicating that the effect is frequency-dependent rather than driven by the total electrical load. Prolonged pacing beyond 12 h led to a decline in cell viability, as evidenced by morphological deterioration of HL-1 cells after 24 h (Figure 5C).

Transfection of HL-1 cells with *Etv1* siRNA prior to field stimulation resulted in a significant reduction of *Kcnk3* mRNA expression under baseline conditions at 1 Hz (*p* = 0.005; *n* = 6; Figure 5D). Remarkably, *Etv1* knockdown via siRNA completely prevented the increase in *Kcnk3* mRNA typically triggered by 5 Hz pacing in control conditions (baseline condition of control group vs. field-stimulation plus *Etv1-*siRNA: *p* = 0.99; *n* = 6).

A similar pattern was observed at the protein level (Figure 5E–F), where 5 Hz field stimulation led to a significant increase in TASK-1 protein expression (*p* = 0.039; *n* = 6), which was fully suppressed by *Etv1* siRNA transfection (baseline condition of control group vs. field-stimulation plus *Etv1-*siRNA: *p* = 0.99; *n* = 6).

Together, these findings demonstrate that ETV1 knockdown prevents pacing-induced upregulation of TASK-1, suggesting a protective role of ETV1 inhibition against tachypacing-induced electrical remodelling.

### ETV1 Binds the *KCNK3* Promoter and Associates with Regulatory Chromatin Features

To investigate whether ETV1 directly binds regulatory regions of the *KCNK3* gene, chromatin immunoprecipitation followed by quantitative PCR (ChIP-qPCR) was performed in HEK-293 cells transiently transfected with human ETV1. Marked enrichment was detected for primer pairs targeting the *KCNK3* promoter region, compared to the negative control (*GAPDH* promoter), and consistent with positive controls targeting known ETV1-responsive promoters (*DUSP6, GPR20*) (Figure 6A).

**Figure 6.**
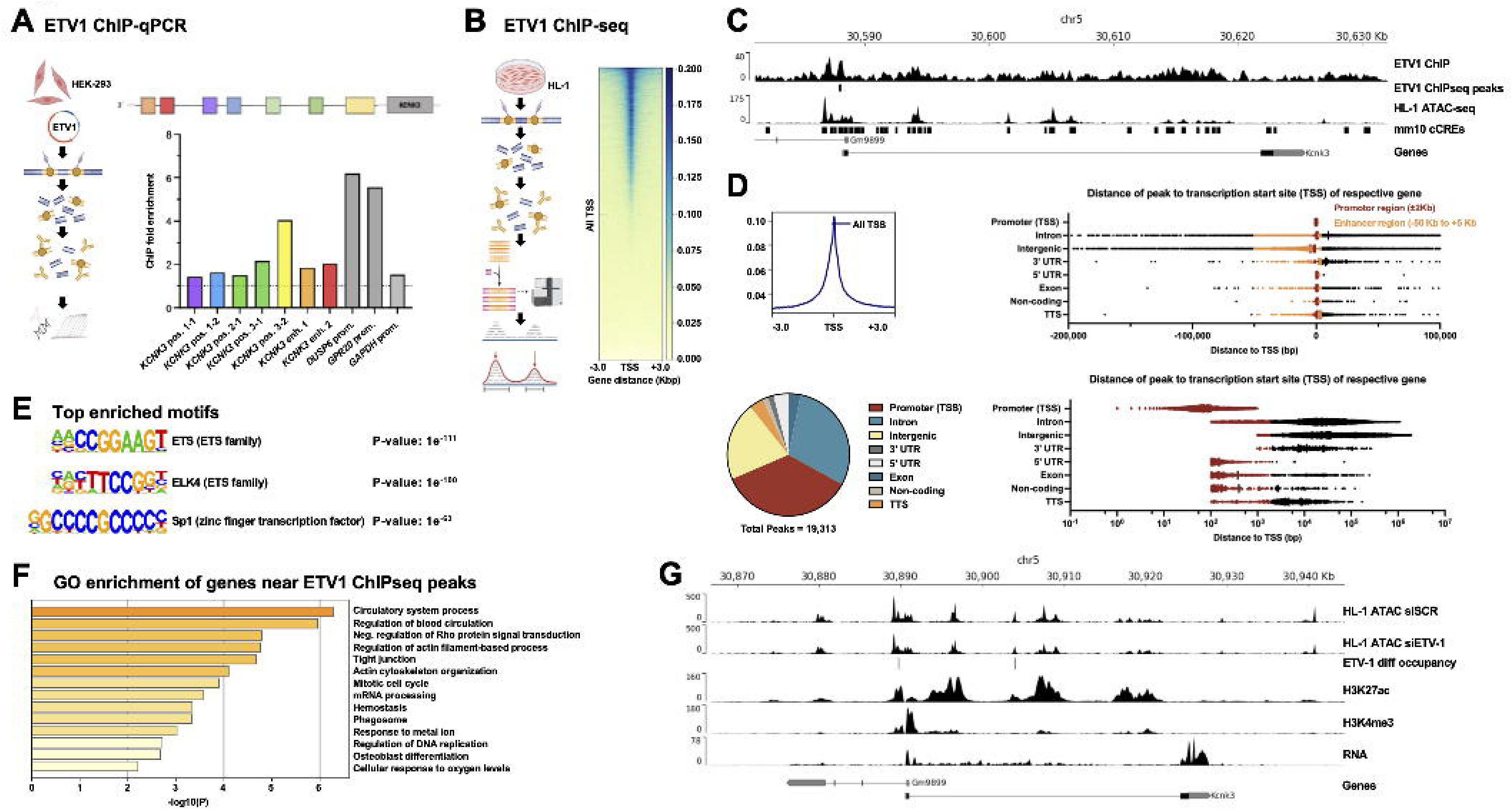
ChIP-based mapping of ETV1 binding sites at the *KCNK3* locus. **A**: Chromatin immunoprecipitation followed by quantitative PCR (ChIP-qPCR) in HEK-293 cells transiently transfected with human ETV1. Fold enrichment values are shown for qPCR primer pairs spanning the enhancer and promoter regions of the *KCNK3* gene. Primer pairs targeting the promoter regions of known ETV1 targets *DUSP6* and *GPR20* serve as positive controls; the *GAPDH* promoter serves as a negative control. **B**: Chromatin immunoprecipitation followed by high-throughput sequencing (ChIP-seq; HL-1 cells) tag density of ETV1 was plotted ±3 kb around annotated transcription start sites (TSS). A heatmap displays normalized read intensity across all TSS, with genes ranked by peak signal. **C**: ChIP-seq signal profile at the *KCNK3* gene locus. A discrete ETV1 peak is located immediately upstream of the transcription start site. Tracks show ETV1 ChIP signal as scaled to input control, ETV1 peak calls, chromatin accessibility (ATAC-seq, HL-1 cells), ENCODE candidate cis-regulatory elements (cCREs) and gene structure. **D**: The average tag profile shows mean enrichment (tags per million) aligned to the TSS (position 0), indicating pronounced promoter-proximal binding. The genome-wide distribution of ETV1 peaks across major genomic features (*bottom right*) shows strongest enrichment at promoters, followed by intronic and intergenic regions, consistent with a transcriptional regulatory role. TTS, transcription termination site; UTR, untranslated region. Distribution of ETV1 ChIP-seq peak distances to the nearest TSS, stratified by genomic feature. Peaks falling within potential promoter regions (±2 kb around the TSS) or putative enhancers (−50 kb to +5 kb) are highlighted in red and orange, respectively. **E**: Known transcription factor motifs enriched in ETV1 ChIP-seq peaks, selected from HOMER analysis based on significance and motif family. Redundant motif variants were excluded for clarity. Each motif is annotated with its transcription factor, DNA-binding domain family, and enrichment *p*-value. **F**: Gene Ontology (GO) enrichment analysis of genes located near ETV1-bound promoter regions. The top enriched biological processes reflect functions related to transcriptional regulation, signal transduction, and cardiac physiology. **G**: ATAC-seq profiles of HL-1 cells treated with scrambled control (siSCR) or siRNA against ETV1 (siETV1) reveal altered chromatin accessibility at the *Kcnk3* promoter upon ETV1 knockdown. ETV1 differential occupancy, histone modification ChIP-seq (H3K27ac, H3K4me3), and RNA-seq are depicted for the *Kcnk3* promoter region.

To globally map ETV1 binding sites, chromatin immunoprecipitation followed by high-throughput sequencing (ChIP-seq) was carried out in HL-1 cells (Figure 6B). Analysis of the genomic distribution of ETV1 binding revealed widespread occupancy across promoter, intronic, and intergenic regions with a strong enrichment at transcription start sites (TSS) (Figure 6C–D). Stratification of peak distances relative to the TSS showed that most promoter-associated peaks clustered within ±2 kb, while distal peaks extended into enhancer-like regions (Figure 6D). At the *KCNK3* locus, a discrete ETV1 ChIP-seq peak was identified directly upstream of the transcription start site, overlapping open chromatin region as determined by assay for transposase-accessible chromatin using sequencing (ATAC-seq) in HL-1 cells. The binding site coincided with candidate cis-regulatory elements (cCREs) and the region amplified in ChIP-qPCR (Figure 6F), supporting direct regulatory engagement of ETV1 at the *KCNK3* promoter.

Motif enrichment analysis of ETV1-bound regions demonstrated significant overrepresentation of ETS-family motifs, including those associated with ETV1 and ETS domain-containing protein Elk-4 (ELK4), as well as distinct motifs from other DNA-binding families, such as the SP1 zinc finger transcription factors Specificity Protein 1 and 2 (SP1, SP2) (Figure 6F and Supplemental Table S6). Gene Ontology analysis of genes located near ETV1 peaks revealed enrichment in biological processes related to transcriptional regulation, signalling, and cardiac function (Figure 6F). Further, a distinct footprint signal was detected at the left enhancer peak, which shows reduced accessibility upon ETV1 knockdown, suggesting additional ETV1 binding outside the main occupancy peak (Figure 6G). This region also coincides with increased H3K27ac and H3K4me3 signal, supporting its role as an active regulatory element, and aligns with elevated *Kcnk3* transcript levels observed in the RNA-seq track.

Finally, a cluster analysis based solely on *ETV1–7* expression levels from RNA-seq data of right atrial appendages in 26 patients revealed distinct molecular subgroups that aligned with the clinical spectrum of atrial cardiomyopathy severity – from no cardiomyopathy through mild and moderate to severe – highlighting the potential involvement of multiple ETV family members in the pathogenesis of atrial remodeling (Figure S6).

## Discussion

The influence of atrial rhythm and left ventricular dysfunction on atrial TASK-1 expression has been well documented in earlier studies ^4,6,8,17,18^. Consistent with previous reports, the results of this study confirm that TASK-1 expression is markedly elevated in atrial tissue from patients with AF compared to those with SR, whereas a stepwise reduction is observed in the context of progressive left ventricular dysfunction. By recapitulating these well-established patterns, our findings not only validate the clinical representativeness of our cohort but also strengthen the evidence implicating TASK-1 as a molecular link, connecting atrial electrical remodelling with both the development of arrhythmogenic substrates and impaired ventricular function.

Interestingly, ETV1 expression patterns mirrored those of TASK-1 in our study. We observed reduced ETV1 expression in patients with impaired LVF, alongside a trend toward increased expression in AF samples. These observations align with recent RNA sequencing data by Rommel *et al.*^14^ which demonstrated a robust upregulation of ETV1 at both the mRNA and protein levels in right atrial tissue samples, derived from AF patients. Furthermore, Yamaguchi et al. (2021), reported diminished ETV1 levels in left atrial tissue from patients with reduced LVF ^19^. However, unlike our study, they did not detect a significant association between ETV1 expression and atrial rhythm status. These conflicting findings may reflect differences in tissue origin (left versus right atrial samples) as well as disparities in patient selection. Notably, the Yamaguchi et al. cohort was restricted to individuals with NYHA class II or higher heart failure, potentially biasing the results toward structural remodelling rather than rhythm-specific effects.

Upon detailed examination of ETV1 and TASK-1 expression profiles in our patient cohort, as well as in an independent RNA-sequencing dataset of 276 human left atrial appendage samples (GEO accession GSE69890^15^) we identified a robust positive correlation between the two transcripts. This association was mechanistically supported by both siRNA-mediated knockdown and pharmacological inhibition of ETV1 using the small molecule inhibitor BRD32048. In HL-1 cardiomyocyte-like cells, suppression of ETV1 consistently led to a marked reduction in TASK-1 expression at both mRNA and protein levels. Importantly, these findings were recapitulated in human atrial fibroblasts and an isolated porcine fibroblast model, highlighting the potential cross-species and cross-cell type relevance of this regulatory axis.

The functional consequences of ETV1 inhibition were further explored using patch-clamp electrophysiology. Both siRNA- and BRD32048-mediated downregulation of ETV1 in HL-1 cells resulted in a significant prolongation of APDLL, without affecting early repolarisation (APDLL) or APA. This selective prolongation is consistent with reduced TASK-1 currents, which predominantly contribute to the terminal repolarisation phase of the atrial AP ^6,8,20^.

Furthermore, indirect patch-clamp measurements with isolation of TASK-1 currents using the experimental antiarrhythmic substance A293 revealed a significant, time-dependent reduction following ETV1 suppression. Given that A293 is a high-affinity TASK-inhibitor ^21–23^, the observed current reduction strongly supports a functional link between ETV1 expression and TASK-1 channel activity. Taken together, these data establish ETV1 as a transcriptional regulator of TASK-1, with direct impact on atrial excitability.

While the functional experiments in HL-1 cardiomyocytes showed a clear reduction in TASK-1 expression and current following ETV1 inhibition, these findings have shown similar tendencies in isolated native human atrial cardiomyocytes from patients based on the limitation of incubation interval with an ETV1 inhibitor. Specifically, A293-sensitive current measurements showed a modest trend towards current reduction in the treatment groups using ETV1 inhibitor-treated cells after 6 hours of incubation. The lack of a robust effect is likely due to the limited duration of drug exposure. Protein turnover rates vary considerably depending on cell type, physiological state and intrinsic properties of the protein itself.

According to the N-end rule proposed by Bachmair *et al.* ^23^, protein stability is largely influenced by the identity of the N-terminal residue. Applying this principle, TASK-1 is predicted to be a highly stable protein with an estimated half-life in excess of 20 hours. It is therefore plausible that a 6 hour incubation period was insufficient to induce measurable degradation or suppression of TASK-1 protein levels in native human cardiomyocytes. A key limitation of the experiments in freshly isolated human cardiomyocytes is, however, the short incubation time, as prolonged treatment beyond 6 hours compromises cell viability and recording quality. Thus, while a trend towards reduced TASK-1 current following ETV1 inhibition was observed, these findings should be interpreted with caution.

Tachypacing of HL-1 cardiomyocytes induced a time-dependent upregulation of TASK-1 expression, peaking at 12 hours, mimicking TASK-1 upregulation in atrial remodelling. To assess the therapeutic potential of ETV1 modulation *in vitro*, HL-1 cells were pretreated with either a pharmacological ETV1 inhibitor or ETV1-targeting siRNA. Both interventions effectively abolished the pacing-induced increase in TASK-1, highlighting a protective effect of ETV1 inhibition against stress-induced electrical remodelling.

Although the tachypacing model in HL-1 cells represents a simplified system and cannot fully replicate the multifactorial and heterocellular nature of AF *in vivo*, it provides a controlled platform to dissect the molecular responses to increased pacing frequency. In this context, our findings provide mechanistic support for the hypothesis that ETV1 knockdown attenuates TASK-1 upregulation under pro-arrhythmic conditions. Although further validation in more physiologically relevant models is warranted, these data highlight the translational potential of targeting ETV1 as a novel strategy to prevent maladaptive atrial electrical remodelling.

To confirm a direct transcriptional link between ETV1 and TASK-1, we performed ChIP-qPCR and genome-wide ChIP-seq. ChIP-qPCR analysis revealed significant enrichment of ETV1 binding at a site close to the *KCNK3* promoter region, strongly supporting direct transcriptional regulation. This was further corroborated by ChIP-seq data, which demonstrated pronounced ETV1 occupancy in close proximity to the *KCNK3* transcription start site. Notably, this region overlapped with candidate cis-regulatory elements and areas of open chromatin identified by ATAC-seq, indicating transcriptional accessibility and regulatory potential. Motif enrichment analysis confirmed overrepresentation of ETS family binding motifs at ETV1-bound loci. In addition to ETS motifs, binding motifs for Sp1, Sp2, and other Ets-domain transcription factors were also enriched at ETV1-bound loci, potentially reflecting co-binding events or common affinity for GC-rich sequences. This suggests that ETV1 might function within a broader regulatory complex influencing atrial gene regulation. Taken together these findings provide compelling evidence that ETV1 directly engages the *KCNK3* promoter in a chromatin context, reinforcing its role as a transcriptional regulator of TASK-1 in atrial cardiomyocytes.

It is important to recognise potential limitations. First, although the expression data from human atrial tissue provide clinical relevance, the cross-sectional nature of the samples precludes causal inference regarding direct regulation of TASK-1 by ETV1. Second, the mechanistic experiments were primarily performed in HL-1 cells and fibroblasts, which, although widely used, do not fully replicate the electrophysiological or transcriptional profile of adult human atrial cardiomyocytes. Third, the short incubation window in freshly isolated human atrial cells limited our ability to observe protein-level changes in response to ETV1 inhibition, likely due to the long half-life of TASK-1 and declining cell viability *ex vivo*. Future studies using translational large animal systems will be essential to validate the therapeutic potential of targeting ETV1 in atrial arrhythmias.

## Conclusion

In summary, our study identifies ETV1 as a key transcriptional regulator of the atrial-specific potassium channel TASK-1, linking it to rhythm- and LVF-dependent electrical remodelling in atrial cardiomyopathy. Through integrative analysis of patient-derived tissue, *in vitro* models, and functional assays, we demonstrate that ETV1 modulates TASK-1 expression and function, and that its inhibition may attenuate stress-induced arrhythmogenic remodelling. These findings establish a mechanistic foundation for targeting ETV1 in AF therapy and provide a rationale for further exploration of TASK-1-based antiarrhythmic strategies.

## Acknowledgments

We thank Anne Grube, Katrin Kupser and Björn Rogatzki for their excellent technical support. The use of AI tools ChatGPT 4.5 (OpenAI, San Francisco, California, USA) and DeepL (DeepL GmbH Cologne, Germany) to improve the readability of the manuscript is acknowledged, with the clarification that they were not employed for experimental design, data evaluation or interpretation. Part of the figures were created utilizing BioRender (Toronto, Canada).

## Funding

This study was supported by the German Research Foundation (DFG) as part of the CRC 1550 (#464424253). M.B., F.W., M.Kr., P.L., A.P., N.F., R.G., and C.S. are members of the CRC 1550 (#464424253), F.W., M.Kr., P.L., A.P., R.G., and C.S. are members of the CRC 1425 (#422681845). Furthermore, this study was supported by research grants from the German Centre for Cardiovascular Research (DZHK) (81X4500125, 81X4500132, 81X4500124, and 81X4500117). From the German Cardiac Society (DGK) (Research Scholarship DGK082018 to F.W.), from the German Heart Foundation / German Foundation of Heart Research (F/15/18 to F.W., F/41/15 to C.S., and Kaltenbach scholarship to A.P.), from Joachim-Herz Foundation (Addon-Fellowship for Interdisciplinary Life Sciences to F.W.), and from the Else-Kröner Fresenius Foundation (EKFS; EKFS Fellowship and Clinician-Scientist professorship to C.S.).

## Disclosure of interest

F.W. and C.S. have filed a patent application for pharmacological TASK-1 inhibition in treatment of atrial arrhythmia. The remaining authors have reported that they have no relation-ships relevant to the content of this paper to disclose.

## Data availability statement

The data that support the findings of this study are available from the corresponding author upon reasonable request.

## Abbreviations

AF: Atrial fibrillation
AP: Action potential
APA: Action potential amplitude
APD: Action potential duration
APD_50_: APD at 50% repolarisation
APD_90_: APD at 90% repolarisation
ATAC-seq: Assay for transposase-accessible chromatin using
cAF: Chronic AF
cCREs: Cis-regulatory elements
ChIP-qPCR: Chromatin immunoprecipitation followed by quantitative PCR
ChIP-seq: Chromatin immunoprecipitation followed by high-throughput sequencing
DTI: Defined Trypsin Inhibitor
ECM: Extracellular matrix
EDTA: Ethylenediaminetetraacetic acid
EGTA: Ethylenebis(oxyethylenenitrilo)tetraacetic acid
ELK4: ETS domain-containing protein Elk-4
ETS: E twenty-six
ETV1: ETS translocation variant 1
FBS: Fetal bovine serum
GEO: Gene Expression Omnibus
Go: Gene Ontology
HEPES: 4-(2-hydroxyethyl)-1-piperazineethanesulfonic acid
K_2P_: Two-pore domain potassium channel
LVF: Left ventricular function
pAF: Paroxysmal AF
PBS: Phosphate-buffered saline
PMSF: Phenylmethanesulfonyl fluoride
RA: Right atrium
RNAseq: RNA sequencing analysis
RMP: Resting membrane potential
RT-qPCR: Real time quantitative polymerase chain reaction
SEM: Standard error of the mean
SR: Sinus rhythm
TEVC: Two-electrode voltage clamp
TSS: Transcription start sites

