## Supplemental Material for "Transcriptional regulation of the TASK-1 potassium channel by ETV1 – Implications for atrial excitability"

^1^ Department of Cardiology and Pneumology, University Medical Center Goettingen, 37075 Goettingen, Germany; DZHK (German Centre for Cardiovascular Research), Partner Site Lower Saxony, 37075 Goettingen, Germany; ^2^ Department of Cardiology, Heidelberg University Hospital, 69120 Heidelberg, Germany; DZHK (German Centre for Cardiovascular Research), Partner Site Heidelberg/Mannheim, Heidelberg University Hospital, 69120 Heidelberg, Germany; ^3^ Institute of Experimental Cardiology, Heidelberg University Hospital, 69115 Heidelberg, Germany; ^4^ Institute for Computational Biomedicine, Bioquant, Faculty of Medicine, Heidelberg University, Heidelberg, Germany; ^5^ Departmant of Cardiac Surgery, Heidelberg University Hospital, 69120 Heidelberg, Germany.

^†^Authors contributed equally

^*^Correspondence:

Prof. Constanze Schmidt, MD, FESC, FEHRA

University Medical Center Goettingen

Department of Cardiology and Pneumology

Robert-Koch-Str. 40

D-37075 Goettingen, Germany

Running title: ETV1 transcriptionally regulates TASK-1 potassium channel expression

**Supplementary Methods section:**

**Study Patients**

All experiments involving human samples were conducted according to the Declaration of Helsinki after approval by the local Ethics Committee of the Medical Faculty at Heidelberg University in Germany (approval number: S-017/2013). Every patient provided written informed consent before sample collection. For patch-clamp experiments right atrial appendage samples were obtained from 5 patients in SR, 4 in pAF, and 3 in cAF undergoing open heart surgery (Table S1). For RT-qPCR experiments samples from 23 patients in SR, 23 in pAF, and 16 in cAF were used (Table S2). RNAseq experiments were performed on samples from 15 patients in SR and 15 in AF (Table S3). Tissue was collected intraoperatively and immediately placed in pre-chilled cardioplegic solution (5 mM NaCl, 50 mM KCl, 6 mM KH2PO4, 25 mM MgSO4, 250 mM taurine, 25 mM MOPS, 20 mM glucose and 30 mM 2,3-butanedion monoxime (pH 7.0)) at 4 °C until processed for cardiomyocyte isolation or fibroblasts culture. Samples intended for total RNA isolation for RT-qPCR or RNAseq, were flash frozen in liquid nitrogen and stored at -80 °C.

**Animal Handling**

All experiments involving German landrace pigs (*Sus scrofa domesticus*) and African clawed frogs (*Xenopus laevis*) were approved by the local Animal Welfare Committee (Regierungspräsidium Karlsruhe, reference numbers G-232/18, G-165/19, G-67/20, G-229/21, and G-45/22). The experiments were carried out according to the guidelines set forth by the U.S. National Institutes of Health (NIH publication No. 86-23), the EU Directive 2010/63/EU, and the German laws for animal protection. Pig hearts were explanted directly after euthanization in deep anaesthesia and sedation. The atrial tissue was stored in pre-chilled cardioplegic solution at 4 °C until cardiomyocyte isolation or fibroblast culture as described ^1^. For harvesting of *X. laevis* oocytes the frogs were anesthetized with a tricaine bath (1 g/l, pH7.5; Pharmaq Ltd., Hamshire, UK) and the ovarian lobes surgically extracted as previously described ^2^. After collagenase (Collagenase D from *Clostridium histolyticum*; Roche diagnostics, Mannheim, Germany) treatment the defolliculated stage V–VI oocytes were manually selected and used for the experiments.

**Chronic Porcine AF Model**

Pigs were premedicated with azaperone (Elanco, Bad Homburg, Germany), midazolam (Hameln Pharma Plus, Hameln, Germany), and ketamine (Zoetis Deutschland, Berlin, Germany), followed by induction of general anaesthesia using propofol (Fresenius Kabi, Bad Homburg, Germany). Analgesia was provided with buprenorphine (Bayer Vital Tiergesundheit, Leverkusen, Germany). Anaesthesia was maintained with isoflurane (Baxter Deutschland GmbH, Heidelberg, Germany). Atrioventricular nodal ablation was then performed, and dual-chamber pacemakers (Abbott Medical, Eschborn, Germany) were implanted. Atrial fibrillation was induced by repeated burst pacing episodes (30 s, 40 Hz), as previously described earlier ^3^. Pacemaker monitoring enabled continuous rhythm surveillance, and burst pacing was suspended upon detection of spontaneous AF. This protocol allowed for the maintenance of AF for 3 weeks. At the study endpoint, animals were euthanized under deep anaesthesia by intravenous injection of potassium chloride, and hearts were harvested during terminal surgery.

**Cardiomyocyte Isolation from Human and Porcine Cardiac Tissue Samples**

Atrial cardiomyocytes were freshly isolated from human and porcine tissue samples as described before ^3^. Briefly, the tissue samples were cut into 1–2 mm³ pieces for enzymatic digestion, rinsed in an ethylenebis(oxyethylenenitrilo)tetraacetic acid (EGTA)-containing solution and afterwards digested in an EGTA and Ca^2+^-free solution with protease type XXIV (Sigma-Aldrich, St. Louis, MO, USA) and collagenase from clostridium histolyticum type V (Sigma-Aldrich, St. Louis, MO, USA). After 30 min the supernatant was removed and the tissue further digested in a protease free solution. The suspension was centrifuged, and the cell pellet resuspended in a storage solution, as reported ^3^.

**Establishing of Atrial Fibroblast Cell Culture**

To establish atrial fibroblast cultures, a portion of freshly obtained atrial tissue was cut into small chunks and transferred into T75 culture flasks filled with cell culture medium consisting of DMEM (Gibco, Waltham, Ma, USA) and Medium 199 (Gibco, Waltham, MA, USA) in a ratio of 4:1 supplemented with 10% fetal bovine serum (FBS; Gibco, Waltham, MA, USA), 100 U/mL penicillin (Gibco, Waltham, MA, USA), and 100 µg/mL streptomycin (Gibco, Waltham, MA, USA). After reaching a confluence of 90–100% the fibroblasts were passaged using 0.05 % Trypsin-EDTA (Gibco, Waltham, MA, USA) and used for experiments only after the first passaging.

**Quantitative Real-Time Polymerase Chain Reaction**

The total RNA fraction was isolated from HL-1 cells, cardiomyocytes, fibroblasts, or right atrial appendage tissue with TRIzol (Invitrogen, Waltham, MA, USA) according to the manufacturer’s instructions. Subsequently, a reverse transcription was performed using the Maxima First Strand cDNA Synthesis Kit for RT-qPCR (Thermo Fisher Scientific, Waltham, MA, USA) following the supplied instructions. The RT-qPCR was performed in 10 µL reaction volumes, run in triplicate, using the TaqMan Fast Universal PCR Master Mix (Thermo Fisher Scientific, Waltham, MA, USA) and TaqMan Assays (Thermo Fisher Scientific, Waltham, MA, USA). The specific assay ID used in this study are listed in Table S4.

**RNA-seq Analysis**

For a poly(A)-enriched bulk RNA sequencing analysis (RNAseq; Illumina HiSeq, 2x150 bp single index, 20–30 M reads /sample) performed by a commercial provider (Genewiz Germany GmbH, Leipzig, Germany) as previously described ^3^, the total RNA fraction of right atrial appendage tissue samples from human patients with SR (n = 15) and AF (n =15) was used as input. For bioinformatics analysis, raw sequencing data underwent initial quality control using MultiQC, followed by alignment with RNA-STAR, read quantification with featureCounts, and differential expression analysis using the DESeq2 algorithm.

For the correlation analysis of ETV1 and TASK-1 mRNA from 276 human patient left atrial appendix samples a publicly available dataset was used (Gene Expression Omnibus (GEO) accession number GSE69890)^4^. Transcript quantification and normalization were performed using the DESeq2 algorithm.

**HEK 293T Cell Culture**

HEK 293T cells were kept in DMEM Medium (Gibco, Waltham, MA, USA) supplemented with 10 % fetal bovine serum (FBS; Gibco, Waltham, MA, USA), 100 U/mL penicillin (Gibco, Waltham, MA, USA), and 100 µg/mL streptomycin (Gibco, Waltham, MA, USA). After the cells reached a confluence of 80 %, they were split in a ratio of 1:12. For experiments cells in passage 10 to 30 were used.

**Molecular Biology and Transfection of HEK 293T Cells**

Plasmid DNA encoding the human ETV1 open reading frame (ETV1_pet28a) was obtained as a gift from Peter Hollenhorst (Addgene plasmid # 131644) and subcloned in the multipurpose expression vector pMax^+^. For transfection a T75 cell culture flask with HEK 293T at around 70 % confluency was used. The cells were transfected with an pMax^+^-ETV1 plasmid using Lipofectamine 3000 (Thermo Fisher Scientific, Waltham, MA, USA) according to the manufacturer’s instructions and incubated for 48 h before use in ChIP-qPCR experiments.

**HL-1 Cell Culture**

HL-1 cells (Sigma-Aldrich, St. Louis, MO, USA) were kept in Claycomb Medium (Sigma-Aldrich, St. Louis, MO, USA) supplemented with 10 % FBS (HL-1 Cell Screened FBS; Sigma-Aldrich, St. Louis, MO, USA), 100 U/mL penicillin (Gibco, Waltham, MA, USA), 100 µg/mL streptomycin (Gibco, Waltham, MA, USA), 2 mM L-glutamine (Gibco, Waltham, MA, USA) and 0.1 mM norepinephrine (Sigma-Aldrich, St. Louis, MO, USA). Before seeding, the cell-dishes were coated with gelatine/fibronectin (Thermo Fisher Scientific, Waltham, MA, USA). After reaching confluency the cells were split in a ratio of 1:3 to 1:4 using 0.05 % Trypsin-EDTA (Gibco, Waltham, MA, USA) and Defined Trypsin Inhibitor (DTI; Gibco, Waltham, MA, USA). For experiments the cells were used with a passage number of 5 to 10.

**siRNA Transfection of HL-1 Cells**

For transfection around 250,000 HL-1 cells per well were seeded into 6-well plates. Two days after seeding, as the cells reached a confluency of about 70 %, the cells were transfected with siRNA targeting ETV1 (ETV1 Silencer Pre-designed siRNA ID: 115583; Ambion, Thermo Fisher Scientific, Waltham, MA, USA) and a scrambled control siRNA (Silencer Negative Control siRNA #1; Ambion, Thermo Fisher Scientific, Waltham, MA, USA) using Lipofectamine 3000 (Thermo Fisher Scientific, Waltham, MA, USA) according to the manufacturer’s instructions and incubated for 48 h before downstream analyses.

**Patch-Clamp Measurements**

Patch-clamp measurements for outward potassium current measurements were performed in voltage-clamp mode with a whole-cell ruptured patch configuration on freshly isolated human cardiomyocytes or HL-1 cells. The pipettes had a resistance of 2–5 MΩ and were backfilled with a solution containing 60 mM KCl, 65 mM potassium glutamate, 5 mM EGTA, 2 mM MgCl_2_, 3 mM K_2_ATP, 0.2 mM NaGTP, and 5 mM HEPES (pH 7.2). The cells were kept in bath solution containing 140 mM NaCl, 5.4 mM KCl, 1 mM MgCl_2_, 1 mM CaCl_2_, 0.3 mM NaH_2_PO_4_, 5 mM HEPES, and 10 mM Glucose (pH 7.4). The cells were clamped at a holding potential of -80 mV and a pulse protocol with pulses of 500 ms starting from -60 mV with an increment of 10 mV until 60 mV were applied. To measure TASK-1 currents the cells were measured at baseline and after application of 200 nm A293. The TASK-1 current was calculated by subtraction of the current after A293 application from the respective baseline current measurement.

Patch-clamp recordings of action potentials from HL-1 cells were performed in current-clamp mode and whole-cell ruptured patch configuration. The pipette solution contained 140 mM KCl, 1 mM MgCl_2_, 5 mM EGTA, 10 mM HEPES, 5 mM K_2_ATP, 3 mM sodium creatine phosphate and 0.1 mM NaGTP (pH 7.2). The bath solution contained 140 mM NaCl, 5 mM KCl, 1 mM MgCl_2_, 1 mM CaCl_2_, 10 mM HEPES, and 10 mM glucose (pH 7.4). The cells were clamped with an average holding current density of -0.9 pA/pF and action potentials were elicited by 10 brief current pulses (5 ms, 500 pA) at 0.5, 1 or 2 Hz stimulation rate every minute. All patch-clamp experiments were conducted using the Axopatch 200B amplifier and Axon Digidata 1550B digitizer (Axon Instruments, Foster City, USA), with data acquisition and analysis performed using pCLAMP software versions 10 and 11 (Axon Instruments, Foster City, USA).

**Two-Electrode Voltage Clamp Measurements**

For two-electrode voltage clamp (TEVC) measurements oocytes from *Xenopus Laevis* were injected with 1.5 ng per oocyte of human TASK-1 cRNA as described elsewhere ^2^. The cells were incubated for 48 h at 18 °C before the measurements were performed. The pipettes had a resistance of 1.5–3 ΩM and were backfilled with 3 M KCl solution. The bath solution contained 101 mM NaCl, 4 mM KCl, 2 mM MgCl_2_, 1.5 mM CaCl_2_, and 10 mM HEPES (pH 7.4). During measurements the cells were clamped at a holding potential of -80 mV. Voltage pulses had a duration of 500 ms starting from -140 mV in increments of 20 mV up to 60 mV (see pulse protocol depicted). Two-electrode voltage-clamp recordings were performed using an OC-725C amplifier (Warner Instruments, Hamden, CT, USA) in combination with a Digidata 1322A or 1550A digitizer (Axon Instruments, Foster City, CA, USA), and data were acquired and analysed using pCLAMP software versions 9 or 10 (Axon Instruments).

**Immunoblot**

The Western Blot analyses was performed as described before ^5^. Briefly, proteins were isolated from HL-1 cells with RIPA-Buffer containing 50 mM TRIS (pH 7.4), 0.5 % NP-40, 0.25 % sodium deoxycholate, 150 mM NaCl, 1 mM EDTA, 1 mM Na_3_VO_4_, 1 mM NaF, and 0.1 % SDS after addition of a cOmplete mini protease inhibitor cocktail (Roche Diagnostics, Mannheim, Germany). The isolated total proteins were quantified using a Pierce BCA protein assay kit (Thermo Fisher Scientific, Waltham, MA, USA) according to the manufacturer’s instructions. The Western Blots were performed in a wet blot setting with primary antibodies against TASK-1 (APC-024; Alomone Labs, Jerusalem, Isreal) and GAPDH (G8140-01; US Biological, Salem, MA, USA) and secondary antibodies Anti-Rabbit (Na934V; Cytiva, Marlborough, MA, USA) and Anti-Mouse (Sc-516102; Santa Cruz Biotechnology, Dallas, TX, USA) respectively. Quantification was done using Fiji ^6^.

***In vitro* Models of Different Cardiomyopathy Subtypes**

For the simulation of different cardiomyopathy subtypes in cell culture, HL-1 cells were seeded in 6-well plates with a density of 250,000 cells per well and incubated until they reached a confluency of 80 %. The cells were than either treated with 200 nM angiotensin II (AdooQ Bioscience, Irvine, CA, USA), 10 µM isoprenaline (Biorbyt, Durham, NC, USA), or 4.5 g/L glucose (DMEM with high Glucose; Gibco, Waltham, MA, USA) and incubated for up to 72 h at 37 °C and 5 % CO_2_ or not treated with any substances and incubated at 37 °C and only 1.0 % O_2_ for up to 24 h. Cells were harvested after 24 h, 48 h and 72 h for mRNA and protein isolation and subsequent analyses for the substance treated groups and after 8 h, 12 h and 24 h for the hypoxia group.

**Rapid Field-Stimulation of HL-1 Cells**

For rapid field-stimulation the cells were seeded in 6-well plates with a density of 250,000 cells per well and incubated until they reached a confluency of 80 %. The cells were paced for up to 24 h with a frequency of 5 Hz as treatment or a frequency of 1 Hz as control (C-Pace EM; Ionoptix, Westwood, MA, USA). After 2 h, 4 h, 8 h, 12 h, and 24 h samples for mRNA and protein isolation were taken and TASK-1 expression analysed. To test the effects of siRNA against *Etv1* on the expression of TASK-1 in a setting of tachypacing the cells were transfected with siRNA against *Etv1* and control siRNA and afterwards paced for 12 h. The cells were harvested for mRNA and protein isolation and subsequent analyses.

**ChIP-qPCR**

For chromatin immunoprecipitation followed by quantitative polymerase chain reaction (ChIP-qPCR) a modified Farnham labs protocol was used ^7^. Following transient Lipofectamin 3000 transfection with pMax^+^ETV1, 20,000,000 HEK-293T cells were fixed with 1 % formaldehyde for 10 min. Chromatin was fragmented by sonication for 10 min (30 s on/ 30 s off) with a Sonifier 250 (Branson Ultrasonics, Brookfield, CT, USA) with a duty cycle of 20 % and an output control of 2. For immunoprecipitation a polyclonal anti-ER81/ETV1 antibody (ab136121; Abcam, Cambridge, United Kingdom) was used. The qPCR was done using SYBR Green mix (PowerSYBR Green PCR Master Mix; AppliedBiosystem, Woolston, Warrington, UK) and individually designed primers (Table S5).

**ChIP-seq**

For chromatin immunoprecipitation followed by DNA sequencing (ChIP-seq) HL-1 cells were fixated for 15 min at room temperature through addition of freshly prepared formaldehyde solution containing 11 % formaldehyde, 0.1 M NaCl, 1 mM ethylenediaminetetraacetic acid (EDTA), and 50 mM 4-(2-hydroxyethyl)-1-piperazineethanesulfonic acid (HEPES) to the cell media. The stop the fixation 1/20 volume of a 2.5 M glycine solution was added for 5 min and the cells scraped form the surface. The cells were first washed with 0.5 % Igepal in phosphate-buffered saline (PBS) and afterwards with PBS-Igepal with the addition of 1mM phenylmethanesulfonyl fluoride (PMSF). After removal of the supernatant the cell pellet was flesh frozen in liquid nitrogen and shipped on dry ice to active motif (Carlsbad, CA, USA). The ETV1 ChIP-seq experiments were performed by a commercial vendor (active motif, Carlsbad, CA, USA) using a polyclonal rabbit anti ETV1 antibody (HPA077249; Sigma-Aldrich, St. Louis, MO, USA).

Sequencing was carried out on an Illumina platform, and raw reads were trimmed using Trim Galore! and assessed for quality with FastQC. Reads were aligned to the mm10 reference genome using BWA and duplicate reads were marked with Picard. Peak calling was conducted using MACS2. Peak annotation and motif enrichment were performed using HOMER, with visualization in IGV and MultiQC. Peaks overlapping putative regulatory regions were intersected with publicly available ENCODE cCRE datasets.

H3K27ac and H3K4me3 ChIP-seq was performed using the Low Cell ChIP Kit (Active Motif, 53086) following the manufacturer’s protocol. Antibodies against H3K27ac (Active Motif, 39133), H3K4me3 (Abcam, GR3213864-23), and H3K4me1 (Active Motif, 39297) were used. Libraries were prepared with the NEBNext Ultra DNA Library Prep Kit for Illumina (New England Biolabs, Frankfurt, Germany) according to the manufacturer’s instructions. Library amplification was monitored in real time using EvaGreen on a qPCR cycler (Agilent) and stopped at the fluorescence inflection point.

**ATAC*-*seq**

The assay for transposase-accessible chromatin using sequencing (ATAC-Seq) experiments were performed as previously reported ^8^. Briefly, per condition 50,000 HL-1 cells were used for the transposase reaction with 2.5 µL Tagment DNA Enzyme 1 (Illumina) followed by a purification with the Qiagen MinElute Reaction Cleanup Kit (Qiagen, Hilden, Germany). The generation and amplification of the libraries was done using the NEBNext High-Fidelity 2x PCR Master Mix (New England Biolabs, Ipswich, MA, USA). AMPure XP Beads (Beckmann Coulter, Brea, CA, USA) were used for cleanup before sequencing on a Novaseq6000 (Illumina) in 75bp PE mode. Adapter sequences were removed from the reads using Trim Galore! (v 0.6.7). Paired-end reads were mapped to the mouse genome (mm10) using Bowtie2 (v 2.5.3) with options '--very-sensitive -X 1000 --dovetail'. Mitochondrial reads were filtered out and PCR duplicates were removed using Picard (v 3.1.1). Data from 3 biological replicates was merged for visualization.

**Statistic**

Data were presented as mean with standard error of the mean (SEM) if not stated otherwise. For testing of statistical significance ordinary one-way ANOVA was used to compare more than two groups with Dunnetts or Šídák’s correction for multiple comparisons as indicated. For comparisons of continuous variables in only two groups unpaired Student’s t-test was used and for categorical variables Fisher’s exact test. p-values < 0.05 were considered statistically significant. For correlation analysis the Pearson correlation coefficients were determined. Statistical analyses were performed using GraphPad Prism version 10.4.1 (GraphPad Software, Boston, MA, USA).

**Supplementary Figures:**

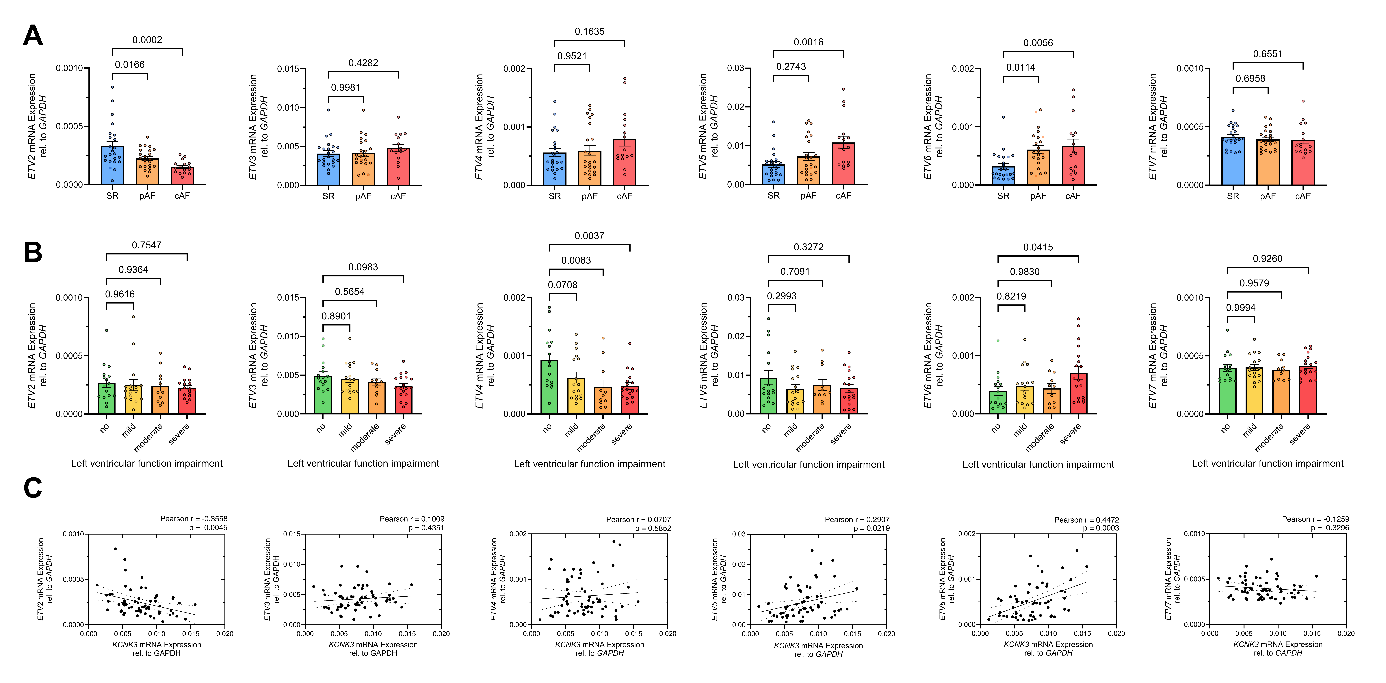

**Figure S1: Expression of *ETV2*–7 stratified by atrial rhythm and degree of left ventricular dysfunction (LVF)** **A–B**: *ETV2–7* mRNA expression relative to *GAPDH* mRNA quantified by quantitative real-time polymerase chain reaction (RT-qPCR) in right atrial appendage tissue samples from patients in sinus rhythm (SR; *n* = 23), paroxysmal atrial fibrillation (pAF, *n* = 23), and chronic atrial fibrillation (cAF; *n* = 16) stratified by rhythm status (A) and LVF impairment (B). **C**: Correlation of *KCNK3* mRNA (encoding for TASK-1) and *ETV2–7* mRNA expression relative to *GAPDH* quantified using RT-qPCR (*n* = 62). Data are presented as mean ± standard error of the mean (SEM). Group differences were assessed using ordinary one-way ANOVA followed by Dunnett’s post hoc correction for multiple comparisons. Statistical significance was defined as *p*-values < 0.05. Pearson correlation coefficients and corresponding *p*-values are indicated within the plots.

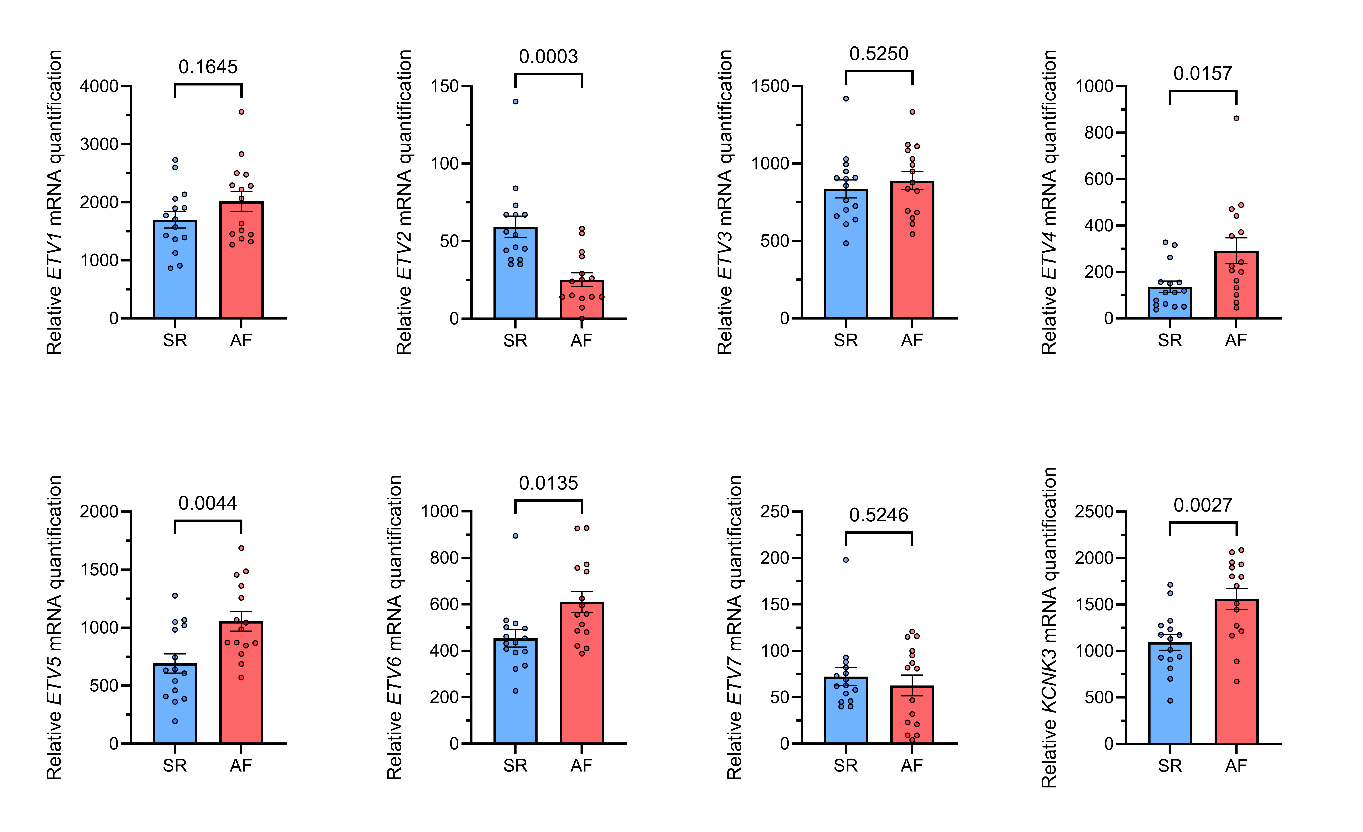

**Figure S2: Relative quantification of *ETV2–7* mRNA levels, stratified by atrial rhythm.** RNAseq expression data of *ETV2*–*7* and *KCNK3* (encoding for TASK-1) in right atrial appendage tissue samples from patients with sinus rhythm (SR; n = 15) and atrial fibrillation (AF; n = 15). Data are presented as mean ± standard error of the mean (SEM). Group comparisons were performed using unpaired Student’s *t*-test and *p*-values < 0.05 were considered statistically significant.

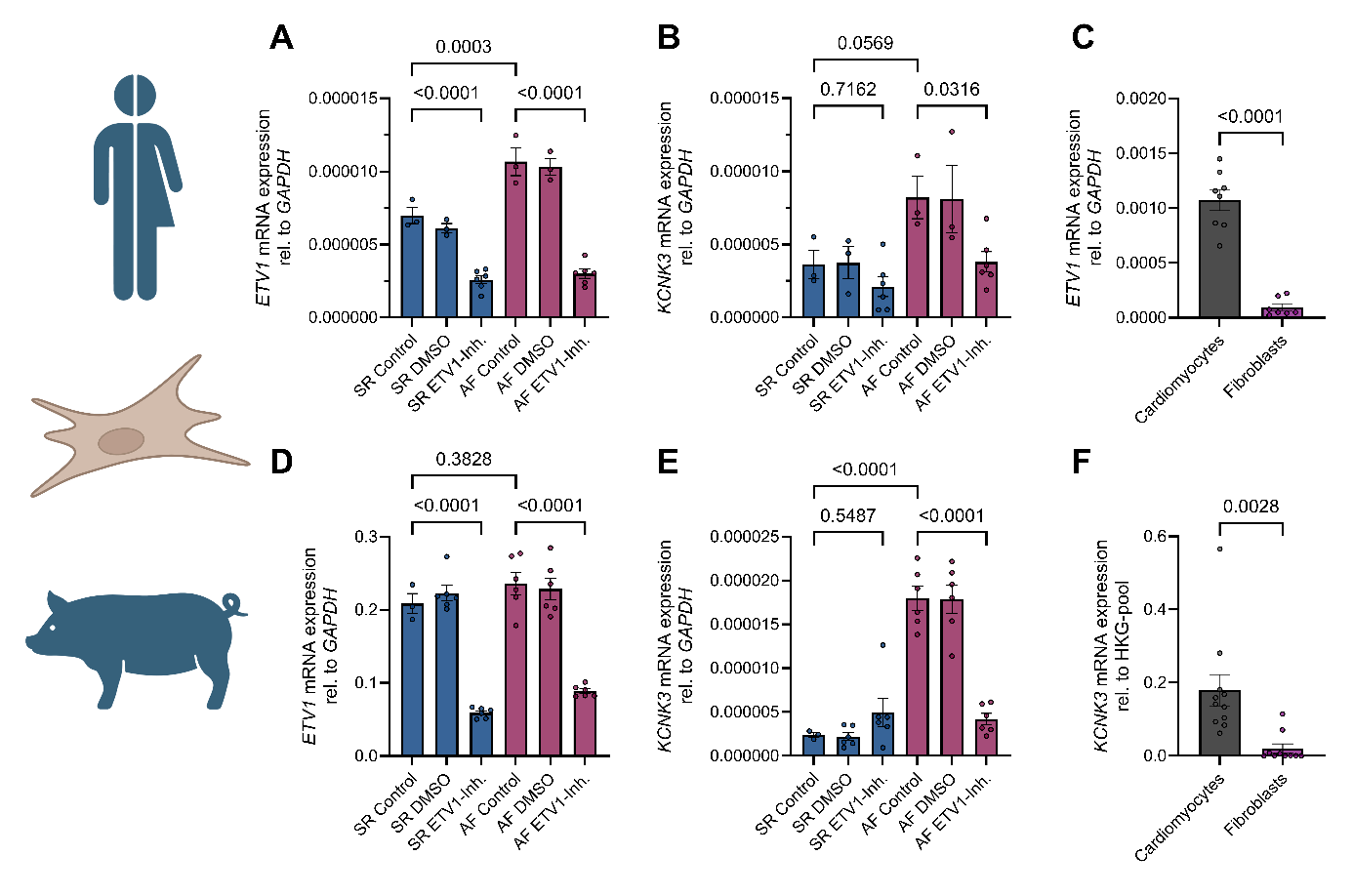

**Figure S3: ETV1 inhibition leads to reduced *KCNK3* mRNA levels in human and porcine fibroblasts. A–B**: *ETV1* mRNA (A, *n* = 3–6) and *KCNK3* mRNA (B, *n* = 3–6) expression relative to *GAPDH* in human atrial fibroblasts, isolated from patients with sinus rhythm (SR; *n* = 3) and atrial fibrillation (AF; *n* = 3) in control conditions (SR Control and AF Control), after treatment with 10 µM of the specific ETV1 inhibitor BRD32048 for 72 h (SR ETV1-Inh. and AF ETV1-Inh.) and corresponding DMSO controls (SR DMSO and AF DMSO). **C**: Comparison of *ETV1* mRNA expression relative to *GAPDH* among human atrial cardiomyocytes (*n* = 8) and fibroblasts (*n* = 7). **D–E**: *ETV1* mRNA (D, *n* = 3–6) and *KCNK3* mRNA (B, *n* = 3–6) levels relative to *GAPDH* in porcine atrial fibroblasts from pigs with SR (*n* = 3) and burst-pacing induced AF (*n* = 3) in control conditions (SR Control and AF Control), after treatment with 10 µM of the specific ETV1 inhibitor BRD32048 for 72 h (SR ETV1-Inh. and AF ETV1-Inh.) and corresponding DMSO controls (SR DMSO and AF DMSO). **F**: Comparison of the *KCNK3* mRNA expression relative to a pool of housekeeping genes (HKG-pool; *GAPDH* and *IPO8*) among human atrial cardiomyocytes (*n* = 10) and fibroblasts (*n* = 11). Please note that data from the subfigure S3F was already published as part of Wiedmann et al., 2022 ^9^. Data are presented as mean ± standard error of the mean (SEM). Ordinary one-way ANOVA with Šídák’s correction for multiple comparisons was used for group comparisons involving more than two groups (A, B, D, E). For comparisons of two groups only, unpaired Student’s *t*-test was applied (C+F). Statistical significance was defined as *p*-values < 0.05.

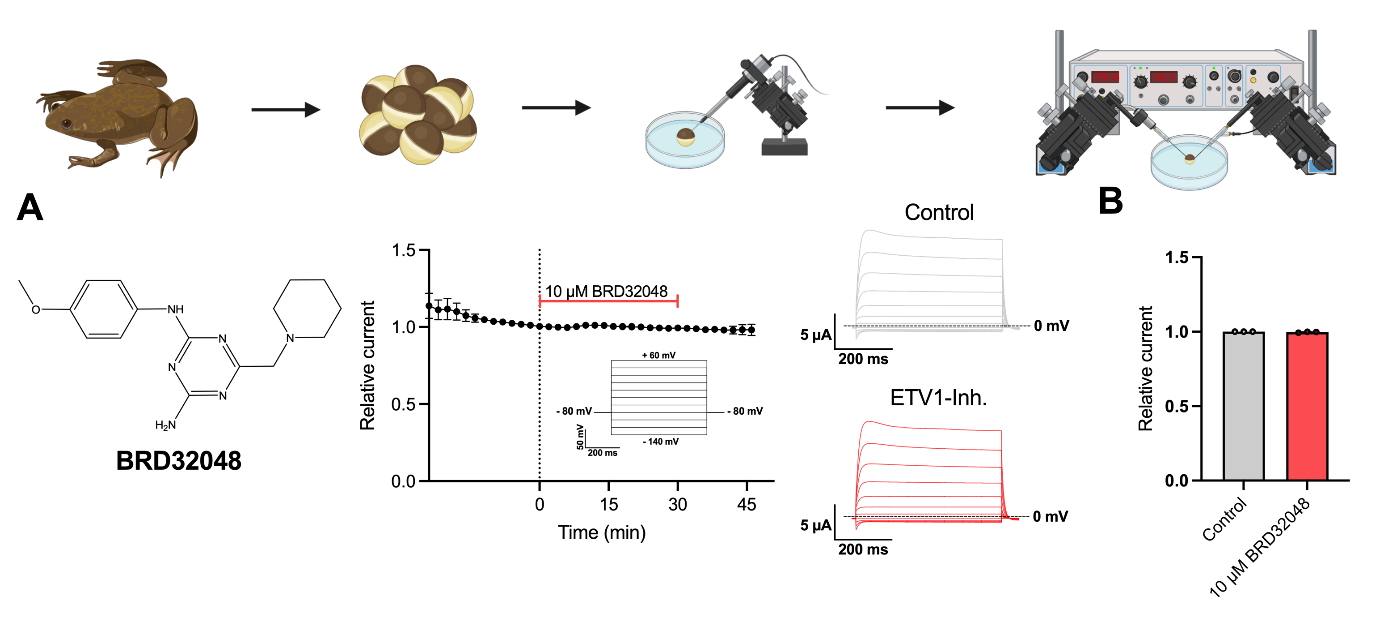
**Figure S4: BRD32048 Does Not Directly Block TASK-1 Channels. A**: TASK-1 current measurements in *Xenopus laevis* oocytes heterologously expressing hK_2P_3.1/TASK-1 channel subunits using the two-electrode voltage clamp (TEVC) technique. Time course of TASK-1 currents during a stabilization period, followed by acute application of 10 µM BRD32048 for 30 min and a 15 min wash-out phase (n = 3) and representative macroscopic TASK-1 current traces indicating that BRD32048 does not exert acute, pore-blocking effects on TASK-1 channels. The voltage protocol used to elicit TASK-1 currents is shown as inset. B: Comparison of the relative current at the end of the stabilization period (Control) and after application of 10 µM BRD32048 for 30 min (n = 3). Data are presented as mean ± standard error of the mean (SEM). Paired Student’s *t*-test was used to compare treatment and control groups and *p*-values < 0.05 were considered as statistically significant.

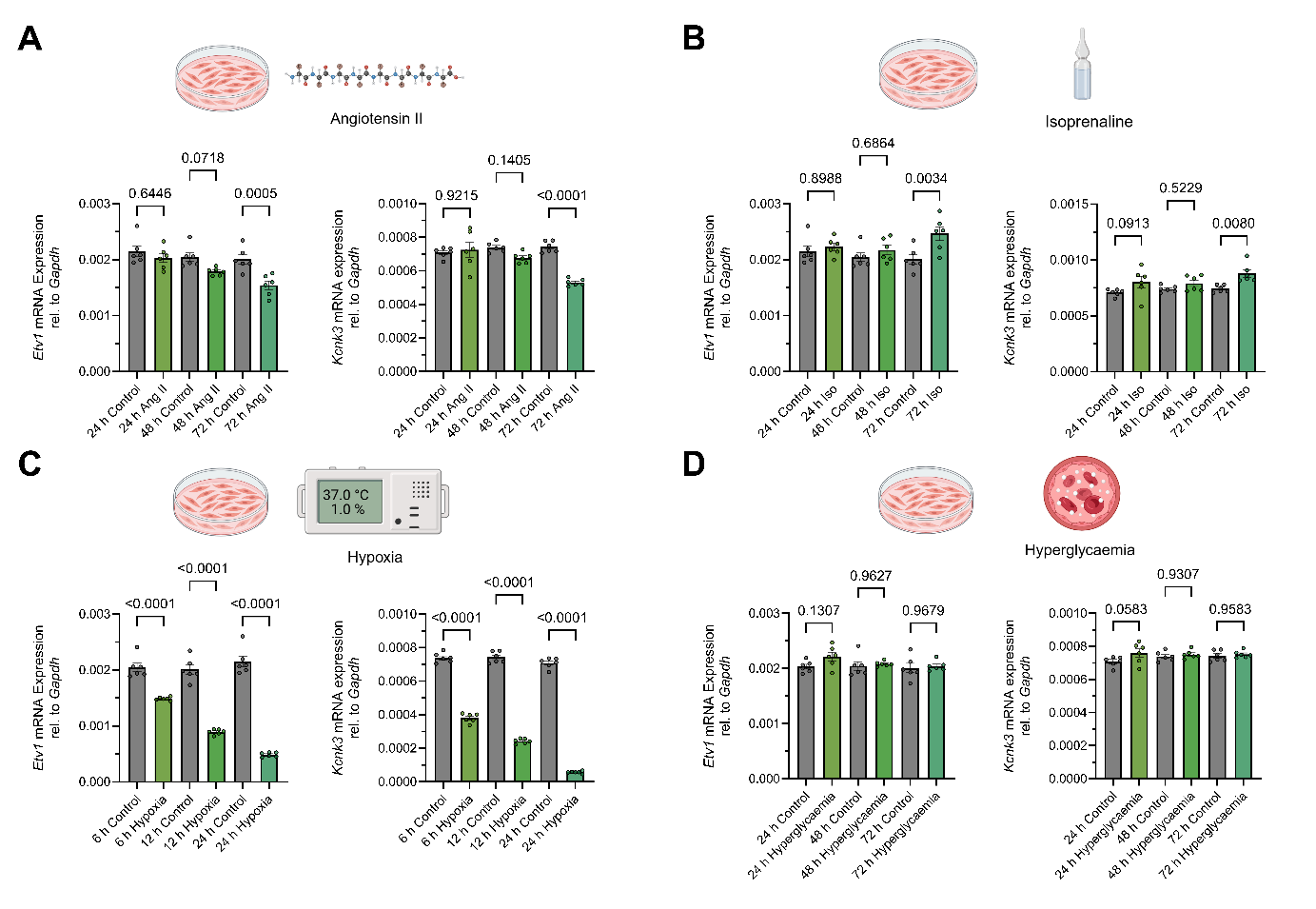
**Figure S5: *Etv1* and *Kcnk3* mRNA levels are changed in different *in vitro* models of atrial cardiomyopathy subtypes.** **A–D**: *Etv1* (n = 6) and *Kcnk3* (n = 6) mRNA expression relative to *Gapdh* mRNA quantified by quantitative real-time polymerase chain reaction (RT-qPCR) in HL-1 cells after treatment with 200 nM angiotensin II (Ang II; A), 10 µM isoprenaline (Iso; B), 1.0% O_2_ (Hypoxia; C), and 4.5 g/L glucose (Hyperglycaemia; D). Data are presented as mean ± standard error of the mean (SEM). Ordinary one-way ANOVA with Šídák’s correction for multiple comparisons was used to compare groups and *p*-values < 0.05 were considered statistically significant.

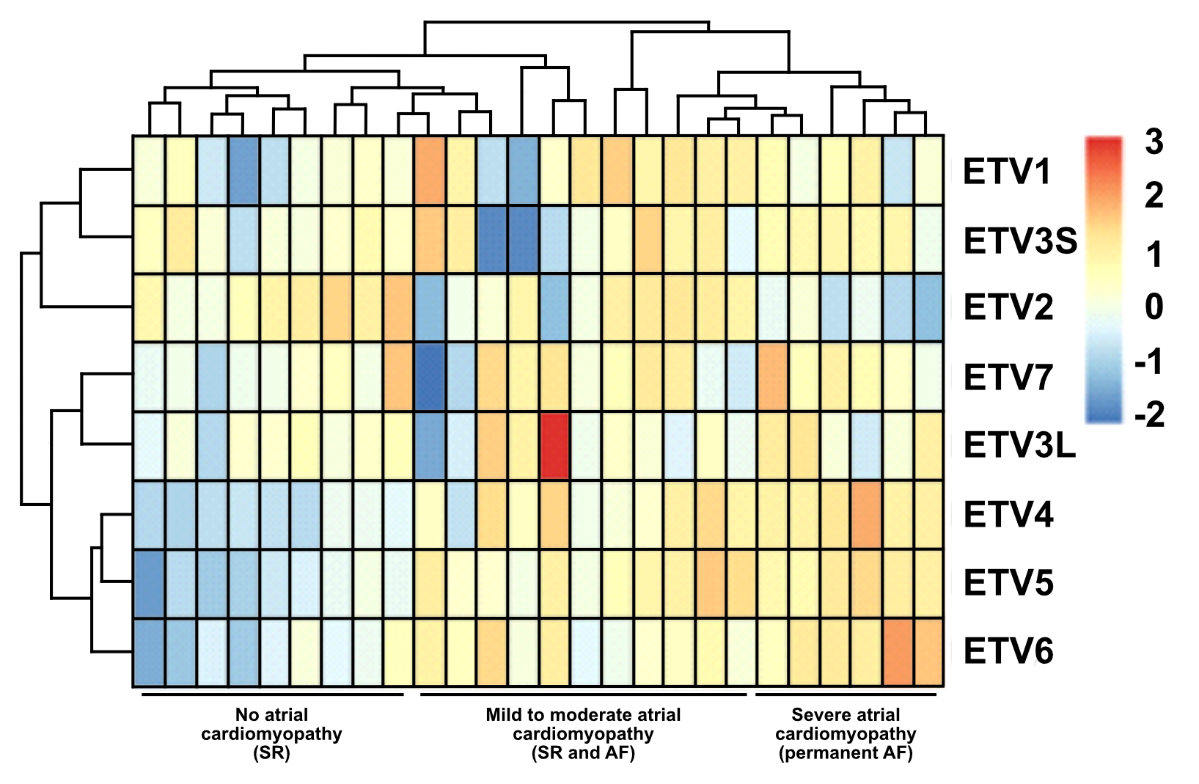

**Figure S6: ETV expression is correlated to cardiomyopathy subtype.** Cluster analysis of relative *ETV1–7* mRNA quantification quantified by RNA sequencing in right atrial appendage samples from 26 human patients with sinus rhythm (SR) and atrial fibrillation (AF). Based on the expression patterns of different ETV family members, patients are arranged along a gradient suggestive of a trajectory of progressive atrial cardiomyopathy, within which distinct regulatory clusters can be delineated.

**Supplementary Tables:**

**Table S1: Baseline characteristics of study patients for patch clamp measurements.** Tissue samples from patients in sinus rhythm (*n* = 5), paroxysmal atrial fibrillation (*n* = 4), and permanent atrial fibrillation (*n* = 3). were used for patch clamp experiments. Unpaired Student’s *t*-test was applied for continuous variables and Fisher’s exact test was used for categorical variables.

AAD: antiarrhythmic drugs; ACE: angiotensin-converting enzyme; AT1: angiotensin II receptor type 1; BMI: body mass index; LVEF: left ventricular ejection fraction; SGLT2: sodium-glucose linked transporter 2.

|  | **Sinus Rhythm**  **(*n* = 5)** | **Paroxysmal Atrial Fibrillation (*n* = 4)** | **Permanent Atrial Fibrillation**  **(*n* = 3)** |
| --- | --- | --- | --- |
| **Demographics** |  |  |  |
| Age, y (mean ± SEM) | 64.6 ± 6.0 | 62.7 ± 16.3 | 67.3 ± 8.3 |
| Female, *n* (%) | 1 (20.0) | 1 (25.0) | 1 (33.3) |
| Height, cm (mean ± SEM) | 178.6 ± 10.5 | 177.5 ± 4.7 | 174.7 ± 10.8 |
| Body weight, kg (mean ± SEM) | 86.4 ± 29.7 | 91.0 ± 19.1 | 80.3 ± 12.7 |
| BMI, kg/m² (mean ± SEM) | 26.8 ± 8.5 | 28.7 ± 5.3 | 26.1 ± 1.2 |
| **Echocardiography** |  |  |  |
| LVEF, % (mean ± SEM) | 53.0 ± 4.7 | 56.5 ± 4.5 | 56.3 ± 5.3 |
| **Medical History, *n* (%)** |  |  |  |
| Hypertension | 4 (80.0) | 4 (100.0) | 1 (33.3) |
| Diabetes | 0 (0.0) | 2 (50.0) | 0 (0.0) |
| Coronary heart disease | 4 (80.0) | 2 (50.0) | 2 (66.7) |
| Hyperlipidaemia | 3 (60.0) | 1 (25.0) | 2 (66.7) |
| **Concomitant medication, *n* (%)** |  |  |  |
| ACE-inhibitors | 3 (60.0) | 1 (25.0) | 1 (33.3) |
| AT1-antagonists | 1 (20.0) | 2 (50.0) | 0 (0.0) |
| Valsartan + sacubitril | 0 (0.0) | 0 (0.0) | 1 (33.3) |
| Statins | 4 (80.0) | 3 (75.0) | 3 (100.0) |
| Digitalis glycosides | 0 (0.0) | 0 (0.0) | 0 (0.0) |
| ß-blockers | 5 (100.0) | 2 (50.0) | 2 (66.7) |
| SGLT2-inhibitors | 0 (0.0) | 2 (50.0) | 0 (0.0) |
| Class III AADs | 0 (0.0) | 0 (0.0) | 0 (0.0) |

**Table S2: Baseline characteristics of study patients for quantitative Real-Time Polymerase Chain Reaction experiments.** Tissue samples from patients in sinus rhythm (*n* = 23), paroxysmal atrial fibrillation (*n* = 23), and permanent atrial fibrillation (*n* = 16). were used for patch clamp experiments. Unpaired Student’s *t*-test was used for continuous variables and Fisher’s exact test for categorical variables, where ** depicts *p* < 0.01.

AAD: antiarrhythmic drugs; ACE: angiotensin-converting enzyme; AT1: angiotensin II receptor type 1; BMI: body mass index; LVEF: left ventricular ejection fraction; SGLT2: sodium-glucose linked transporter 2.

|  | **Sinus Rhythm**  **(*n* = 23)** | **Paroxysmal Atrial Fibrillation (*n* = 23)** | **Permanent Atrial Fibrillation**  **(*n* = 16)** |
| --- | --- | --- | --- |
| **Demographics** |  |  |  |
| Age, y (mean ± SEM) | 67.5 ± 11.5 | 69.8 ± 10.2 | 70.2 ± 9.7 |
| Female, *n* (%) | 5 (21.7) | 9 (39.1) | 4 (25.0) |
| Height, cm (mean ± SEM) | 173.9 ± 7.5 | 172.0 ± 8.4 | 176.2 ± 9.2 |
| Body weight, kg (mean ± SEM) | 86.3 ± 13.6 | 83.1 ± 22.7 | 92.5 ± 19.6 ****** |
| BMI, kg/m² (mean ± SEM) | 28.6 ± 4.4 | 28.2 ± 6.9 | 29.8 ± 5.3 |
| **Echocardiography** |  |  |  |
| LVEF, % (mean ± SEM) | 36.9 ± 11.9 | 40.5 ± 10.6 | 37.3 ± 13.8 |
| **Medical History, *n* (%)** |  |  |  |
| Hypertension | 23 (100.0) | 22 (95.6.0) | 14 (87.5) |
| Diabetes | 6 (26.1) | 10 (43.4) | 6 (37.5) |
| Coronary heart disease | 11 (47.8) | 11 (47.8) | 6 (37.5) |
| **Concomitant medication, *n* (%)** |  |  |  |
| ACE-inhibitors | 9 (39.1) | 11 (47.8) | 9 (56.3) |
| AT1-antagonists | 4 (17.4) | 6 (26.1) | 4 (25.0) |
| Valsartan + sacubitril | 0 (0.0) | 0 (0.0) | 0 (0.0) |
| Statins | 17 (73.9) | 18 (78.2) | 10 (62.5) |
| Digitalis glycosides | 1 (4.3) | 1 (4.3) | 2 (12.5) |
| SGLT2-inhibitors | 0 (0.0) | 0 (0.0) | 0 (0.0) |
| Class III AADs | 0 (0.0) | 1 (4.3) | 2 (12.5) |

**Table S3: Baseline characteristics of study patients for RNAseq experiments.** Tissue samples from patients in sinus rhythm (*n* = 15) and atrial fibrillation (*n* = 15). were used for patch clamp experiments. Unpaired Student’s *t*-test was used for continuous variables and Fisher’s exact test for categorical variables.

AAD: antiarrhythmic drugs; ACE: angiotensin-converting enzyme; AT1: angiotensin II receptor type 1; BMI: body mass index; LVEF: left ventricular ejection fraction; SGLT2: sodium-glucose linked transporter 2.

|  | **Sinus Rhythm**  **(*n* = 15)** | **Atrial Fibrillation**  **(*n* = 15)** |
| --- | --- | --- |
| **Demographics** |  |  |
| Age, y (mean ± SEM) | 69.3 ± 9.1 | 69.9 ± 9.4 |
| Female, *n* (%) | 3 (20.0) | 3 (20.0) |
| Height, cm (mean ± SEM) | 173.7 ± 5.9 | 176.9 ± 11.5 |
| Body weight, kg (mean ± SEM) | 83.4 ± 14.6 | 94.2 ± 13.1 |
| BMI, kg/m² (mean ± SEM) | 27.6 ± 4.6 | 30.2 ± 3.6 |
| **Echocardiography** |  |  |
| LVEF, % (mean ± SEM) | 44.7 ± 15.1 | 40.5 ± 13.6 |
| **Medical History, *n* (%)** |  |  |
| Hypertension | 13 (86.7) | 13 (86.7) |
| Diabetes | 4 (26.7) | 5 (33.3) |
| Coronary heart disease | 14 (93.3) | 15 (100) |
| Hyperlipidemia | 11 (73.3) | 10 (66.7) |
| **Concomitant medication, *n* (%)** |  |  |
| ACE-inhibitors | 4 (26.7) | 8 (53.3) |
| AT1-antagonists | 3 (20.0) | 3 (20.0) |
| Valsartan + sacubitril | 0 (0.0) | 1 (33.3) |
| Statins | 11 (73.3) | 12 (80.0) |
| Digitalis glycosides | 1 (6.7) | 1 (6.7) |
| ß-blockers | 9 (60.0) | 12 (80.0) |
| SGLT2-inhibitors | 0 (0.0) | 0 (0.0) |
| Class I/III AADs | 2 (13.3) | 3 (20.0) |

**Table S4: TaqMan assays used for quantitative Real-Time Polymerase Chain Reaction (RT-qPCR).**

| **Target** | **Species** | **TaqMan Assay** |
| --- | --- | --- |
| *ETV1* | Human | Hs0095191_m1 |
| *Etv1* | Murine | Mm0514804_m1 |
| *ETV1* | Porcine | Ss06866942_m1 |
| *ETV2* | Human | Hs01012852_g1 |
| *ETV3* | Human | Hs01379808_m1 |
| *ETV4* | Human | Hs_00383361_g1 |
| *ETV5* | Human | Hs_00927557_m1 |
| *ETV6* | Human | Hs_00231101_m1 |
| *ETV7* | Human | Hs_00903229_m1 |
| *KCNK3* | Human | Hs_00605529_m1 |
| *Kcnk3* | Murine | Mm_00807036_m1 |
| *GAPDH* | Human | Hs_99999905_m1 |
| *Gapdh* | Murine | Mm_99999915_g1 |

**Table S5: Primers used for chromatin immunoprecipitation followed by quantitative polymerase chain reaction (ChIP-qPCR).**

| **Name** | **Sequence** |
| --- | --- |
| *KCNK3*-Position1-1 (Forward) | CGGGGAGAGTAGCATGAGTG |
| *KCNK3*-Position1-1 (Reverse) | CTGCAGCACCCTATCTAAGCA |
| *KCNK3*-Position1-2 (Forward) | TAAGGTGCAAAGTACTGCGGG |
| *KCNK3*-Position1-2 (Reverse) | CATATCCATCTTCCCCACTGGC |
| *KCNK3*-Position2-1 (Forward) | GGGTGTCATTGGAGGAGTGTAG |
| *KCNK3*-Position2-1 (Reverse) | CCTGACTGAGAGCAGACTCA |
| *KCNK3*-Position3-1 (Forward) | GCACACATTGCCTGAGAACA |
| *KCNK3*-Position3-1 (Reverse) | TTCCTCCCTTGGGCCAGACTC |
| *KCNK3*-Position3-2 (Forward) | CTGAGAACAGGGCTTTCCCTT |
| *KCNK3*-Position3-2 (Reverse) | CTCCTTCCTCCCTTGGGCCAG |
| *KCNK3*-Enhancer-1 (Forward) | AAGTCACTTGGCATCACCCA |
| *KCNK3*-Enhancer-1 (Reverse) | TGGGGGCCAGAGCTATCAAT |
| *KCNK3*-Enhancer-2 (Forward) | ACAGCCTTCCTCTCTGAGCTA |
| *KCNK3*-Enhancer-2 (Reverse) | TGACTCCCAAGTGTGTGCTT |
| *DUSP6*-Promoter (Forward) | GCCCGCTGTTGCAGCTTGTT |
| *DUSP6*-Promoter (Reverse) | GCCGGCTGGAACAGGTTGTG |
| *GPR20*-Enhancer (Forward) | CCCTCCCAGGCTCTCCCCAC |
| *GPR20*-Enhancer (Reverse) | TCCGGGCCTGCTCTCTGTCC |
| *GAPDH*-Promoter (Forward) | TCCCAAAGTCCTCCTGTTTCA |
| *GAPDH* -Promoter (Reverse) | CAGCAGGACACTAGGGAGTCAA |

**Table S6: Known transcription factor motifs enriched in ETV1 ChIP-seq peaks (from HOMER analysis)**

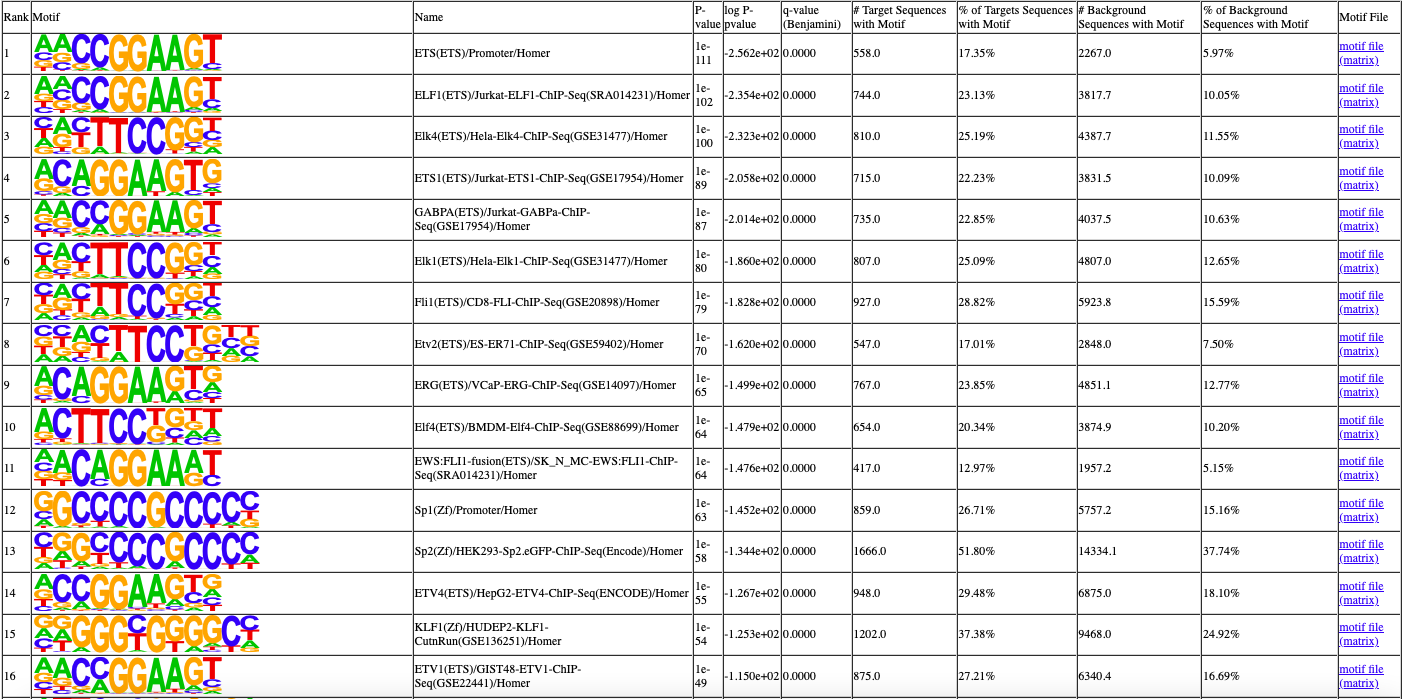
